# The nuclear egress complex of MHV-68 is not essential for nuclear egress but mediates C-capsid specificity

**DOI:** 10.1101/2025.04.10.648238

**Authors:** Saskia Sanders, Carola Schneider, Timothy K. Soh, Elena Kotova, Beatrix Steer, Rudolph Reimer, Zsolt Ruzsics, Heiko Adler, Jens B. Bosse

**Author notes:** To whom correspondence should be addressed.; (J.B.B.); (H.A.). These authors contributed equally to this work (Heiko Adler, Jens B. Bosse).

## Abstract

Herpesvirus capsids must exit the nucleus to undergo additional maturation steps in the cytoplasm, such as secondary envelopment. This process is orchestrated by the nuclear egress complex (NEC), a conserved heterodimer that deforms the inner nuclear membrane and facilitates capsid egress. While NEC proteins are critical for the release of capsids into the cytoplasm and, therefore, efficient replication in ɑ-, β-, and γ-herpesviruses, residual infectivity has been observed in NEC-deficient mutants of certain herpesviruses, including the ɑ-herpesvirus Pseudorabies virus (PrV).

To investigate the role of the NEC in murine gammaherpesvirus 68 (MHV-68), a mutant virus lacking most of the nucleoplasmic NEC component ORF69 (ΔORF69) was generated. Unlike previous studies relying on transfection-based systems, this study demonstrates that NEC-deficient MHV-68 remains replication-competent, albeit with a significant replication defect. Despite the absence of ORF69, the mutant virus produced infectious progeny in non-complementing cells. Quantitative electron microscopy revealed regular nuclear capsid assembly and DNA packaging. However, while the parental virus primarily exported mature, genome-filled C-capsids, the ΔORF69 mutant exhibited cytoplasmic and extracellular emergence of all capsid forms.

These findings indicate that the MHV-68 NEC functions as a quality control checkpoint, selectively facilitating the nuclear egress of mature capsids. In the absence of ORF69, some capsids still egress, likely through alternative, less efficient pathways. Inhibition of cyclin-dependent kinases reduced parental and mutant virus spread, suggesting that γ-herpesviruses exploit host-dependent mechanisms for an alternative nuclear egress route.

**Importance:** Gammaherpesviruses, such as Epstein-Barr virus and Kaposi’s sarcoma-associated herpesvirus, establish lifelong infections and can contribute to cancer. A crucial step in the viral life cycle depends on the nuclear egress of capsids, a process mediated by the viral nuclear egress complex (NEC). Surprisingly, murine gammaherpesvirus 68 (MHV-68) can translocate capsids from the nucleus to the cytoplasm even when a key NEC component, nucleoplasmic ORF69, is disrupted. Deleting most of ORF69 reduced viral replication but did not entirely block nuclear egress. Instead, the mutant virus allowed both mature and immature capsids to leave the nucleus, suggesting that the NEC functions as a quality control checkpoint for mature capsids. Furthermore, host cell kinases seem to enable alternative routes for egress that do not rely on the NEC. These findings further support the notion that NEC is not strictly necessary and suggest the potential exploitation of host-virus interactions for antiviral approaches.

## Introduction

Herpesviruses are a widely distributed family of dsDNA viruses. The Herpesviridae family is divided into three subfamilies: ɑ-, β-, and γ-herpesvirinae, with human pathogenic and highly species-specific γ-herpesviruses such as Epstein-Barr Virus (EBV) and Kaposi’s Sarcoma-Associated Herpesvirus (KSHV) being particularly significant due to their oncogenic potential (1). One hallmark of herpesvirus infections is the ability to establish lifelong latency after primary infection and reactivation under conditions of immunosuppression (1). Murine gammaherpesvirus 68 (MHV-68) serves as a well-established model to study γ-herpesviruses due to its wider host range and primarily lytic life cycle in contrast to the predominantly latent EBV and KSHV in cell culture (1,2).

The virion structure of herpesviruses is highly conserved, comprising a double-stranded DNA genome, a 120–130 nm icosahedral capsid, a protein-rich, amorphous layer called tegument, and an envelope containing glycoproteins essential for host cell entry. Following viral entry, capsids are transported to the host cell nucleus, where viral DNA is injected through nuclear pores to initiate replication and capsid assembly. Packaged capsids containing viral dsDNA are termed C-capsids. In addition to these infectious precursors, two abortive capsid forms, B-capsids, which retain the scaffold, and A-capsids, which lack scaffold and DNA, are present in the nucleus (3,4). Only C-capsids are capable of productive infection and must translocate to the cytoplasm to complete further maturation, requiring an efficient nuclear egress process. The capsid buds at the inner nuclear membrane (INM), where primary envelopment occurs, subsequently fuses with the outer nuclear membrane (ONM) and is then released into the cytoplasm (1).

This process relies on the nuclear egress complex (NEC), a conserved heterodimer composed of a tail-anchored membrane protein and its nucleoplasmic partner, which interact by a conserved hook-groove interaction (5). The NEC has been extensively characterized in α-herpesviruses, such as HSV-1 and Pseudorabiesvirus (PrV) (6), where these proteins correspond to UL34 and UL31, respectively. Similarly, the NEC of β-herpesviruses, especially the human cytomegalovirus (HCMV), has been well studied (7–9). However, among the herpesvirus subfamilies, the NEC of γ-herpesviruses remains the least characterized, leaving critical gaps in our understanding of its function. Since NEC-mediated nuclear remodeling and capsid egress depend on its oligomerization and membrane interactions, structural insights are essential for dissecting its role.

Classical structural biology techniques have laid the foundation for understanding NEC architecture across herpesviruses. X-ray crystallography has provided high-resolution structures of various NEC heterodimers (10–16), while cryo-electron microscopy (cryo-EM) and electron cryo-tomography (cryo-ET) have advanced our understanding of NEC higher-order assemblies, particularly in the context of membrane remodeling, a crucial process in herpesvirus egress (12,17–19). However, these techniques remain highly time- and labor-intensive and have not been applied comprehensively to all herpesviruses, particularly γ-herpesviruses. In this context, AlphaFold is a powerful complementary tool, offering rapid and reliable structural predictions (20). AlphaFold provides valuable insights that guide experimental design by identifying conserved NEC structural features across herpesviruses. Given its demonstrated accuracy in predicting general protein structures (21,22) and viral proteins (23–25), AlphaFold accelerates the investigation of γ-herpesviruses NEC heterodimer and oligomeric structures, bridging the existing knowledge gap.

While structural predictions help refine our understanding of NEC-mediated nuclear egress, herpesviruses may also utilize alternative routes for nuclear egress. In α-herpesviruses, nuclear envelope breakdown (NEBD) has been observed, and nuclear pore dilation has been suggested as an additional route for nuclear egress (26,27). However, in γ-herpesviruses, the molecular mechanisms of nuclear egress remain less well-characterized. The core NEC proteins in EBV have been identified as the tail-anchored membrane protein BFRF1 (28) and nucleoplasmic partner BFLF2 (29), while KSHV encodes homologous proteins ORF67 and ORF69 (30). Similarly, MHV-68, whose genome is colinear with KSHV, encodes ORF67 and ORF69 as NEC components (31,32). Yet, bacterial artificial chromosome (BAC) transfection studies suggest that these proteins are not absolutely essential for viral replication, as viral genomes were still detected at reduced levels (31). This finding raises the possibility that γ-herpesviruses may also employ alternative egress routes, though this mechanism has not yet been characterized.

To address this, we use MHV-68 as a model to investigate the role of the NEC in γ-herpesvirus nuclear egress, focusing on the nucleoplasmic NEC protein ORF69. Specifically, we use AlphaFold to model a partial ORF69 deletion mutant that retains the conserved hook and amino acids required for interaction with ORF67 but lacks the C-term and, therefore, the ability to oligomerize. With quantitative thin-section electron microscopy, we show that the WT NEC acts as a C-capsid-specific quality control checkpoint as mainly C-capsid egress. In ORF69 mutant, capsids accumulate in the nucleus, yet packaging remains unaffected. Despite the absence of a functional NEC, low-level spread is detectable, suggesting an alternative, inherent egress route likely to be CDK pathway dependent. By integrating structural modeling, ultrastructural analysis, and functional assays, this study provides new insights into the NEC’s role in γ-herpesvirus nuclear egress and potential alternative egress mechanisms.

## Results

### Design of MHV-68 ΔORF69

So far, it has been described for KSHV and MHV-68 that the NEC consists of ORF69 and ORF67 (33,34), but the lack of fluorescent mutants makes it challenging to study their function, especially in living cells. We wanted to study the NEC in lytic infection of the γ-herpesvirus MHV-68 and constructed an ORF69 mutant based on the small capsid protein-tagged fluorescent mutant (35). In this study, the parent virus, MHV-68 ORF65mCherry, is hereafter referred to as WT, while the ORF69 deletion mutant, MHV-68 ORF65mCherry ΔORF69, is referred to as ΔORF69. To minimally disrupt the genetic context of ORF69, we used BAC mutagenesis to delete the third in-frame ATG at codon 69 by substituting the A with a T and removing the subsequent TG. The mutation causes a frameshift, leading to a sequence of 40 altered amino acids beginning at position 69 while preserving the wild-type sequence up to amino acid 68. As a result, only the N-term of ORF69 is correctly expressed. Further, this is important because *in silico* monomer predictions of ORF67 suggested that complete deletion of ORF69 could lead to the misfolding of ORF67, and we therefore preserved the N-terminal region of ORF69.

Additionally, a previous study identified three critical residues, F52, F65, and L66, for mediating the ORF67-ORF69 interaction. In addition, the amino acid E68, combined with the three others, forms a group of conserved amino acids on the interaction surface in MHV-68 ORF69 and its homologs in KSHV and EBV (34). Also, the N-term of ORF69 contains highly basic clusters, as seen for other herpesviruses with different proposed functions (36,37). Therefore, our design preserved the ORF69 N-terminus up to amino acid E68 (Figure 1A). BAC digestion with EcoRI, HindIII, and SacI confirmed the mutation compared with WT through additional bands (Figure S1).

**Figure 1:**
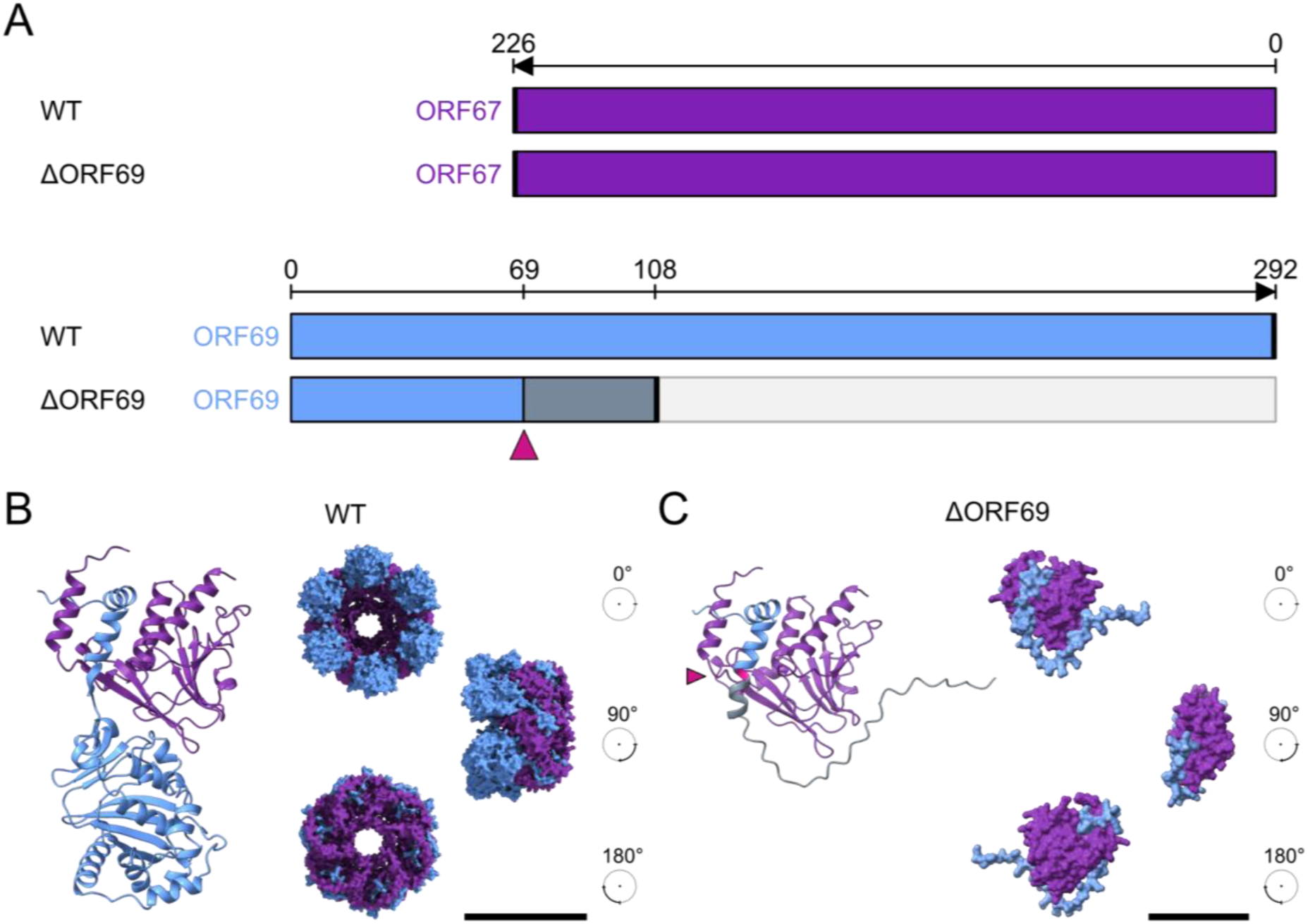
Design of MHV-68 ΔORF69 mutant. (A) Schematic illustration of the NEC proteins of WT and ΔORF69 mutant. The ΔORF69 mutant was generated by deleting the third in-frame ATG codon of ORF69 at amino acid 69 (pink arrowhead), introducing a frameshift mutation leading to a premature stop codon at amino acid 108 (thick black line). ORF69 is shown in blue, ORF67 in purple, the frameshifted region in blue-gray, and the non-expressed downstream region in light gray. (B) AlphaFold3-predicted ORF67-ORF69 heterodimer (ipTM = 0.75, pTM = 0.72) is shown alongside three surface representations of a hexameric ORF67-ORF69 complex, rotated by 90°. Scale bar: 100 Å (C) AlphaFold3-predicted ORF67-ΔORF69 heterodimer (ipTM = 0.74, pTM = 0.65) is shown with three surface representations rotated by 90°. Scale bar: 50 Å.

**Figure S1:**
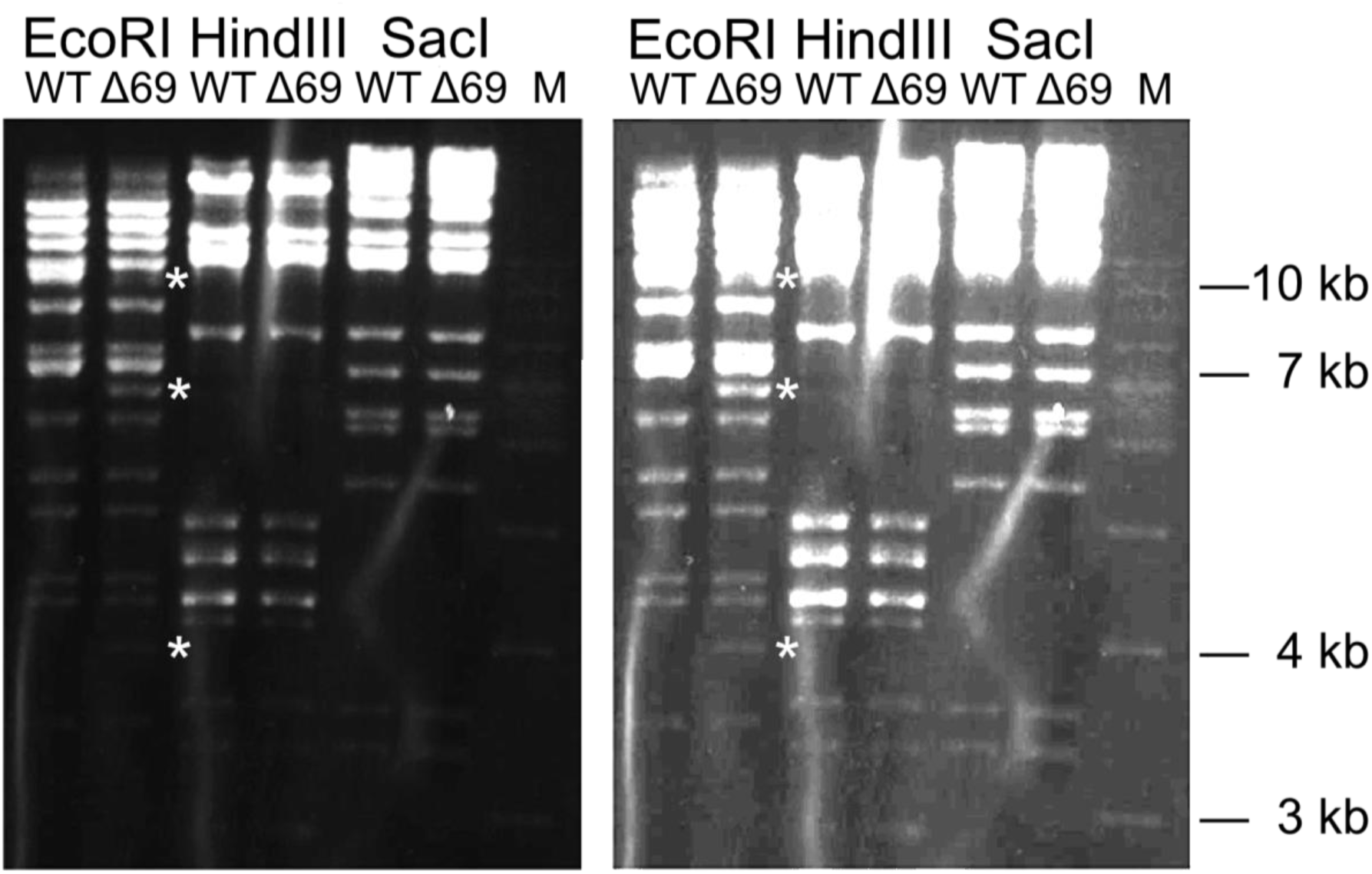
Restriction digest analysis of WT and ΔORF69 MHV-68 BACs. BACs of WT and ΔORF69 were analyzed by digestion with EcoRI, HindIII, and SacI. The left panel shows a low-exposure image, while the right panel presents a high-exposure version. Differences in digestion patterns are indicated by asterisks.

We used the AlphaFold Server (38) to model heterodimers of ORF67-ORF69 (Figure 1B, left) and ORF67-ΔORF69 (Figure 1C, left), which had high confidence scores and are likely interacting. The ORF67-ORF69 heterodimer likely oligomerizes into higher-order up to heterohexameric complexes (Figure 1B). In contrast, the ORF67-ΔORF69 complex remains heterodimeric and does not form higher-order oligomers (Figure 1C). Although the AlphaFold quality scores for ORF67-ORF69 and ORF67-ΔORF69 heterodimers are high, higher order-multimers differ drastically between the WT and the KO mutant (Figure S2). Higher-order multimers of ORF67-ΔORF69 exhibit low confidence scores, indicating limited oligomerization potential, with scores decreasing further as stoichiometry increases.

In contrast, ORF67-ORF69 NEC multimers maintain high confidence scores up to the trimeric form. A notable drop in confidence is observed at the tetramer level, followed by an increase for pentamer and hexamer formations. Additionally, we utilized AlphaFold to model the γ-herpesviruses EBV and KSHV NEC multimers. KSHV consistently achieves high-quality scores in both heterodimer and higher-order multimer forms. Although crystallographic data suggests a dimeric NEC conformation for EBV, our predictions indicate that EBV multimers consisting of three, five, and six heterodimers have higher confidence values than the heterodimeric form (39). These trends are also reflected in the PAE plots and the pairwise comparison of ipTM scores across the proteins (Figure S2B, S2C).

**Figure S2:**
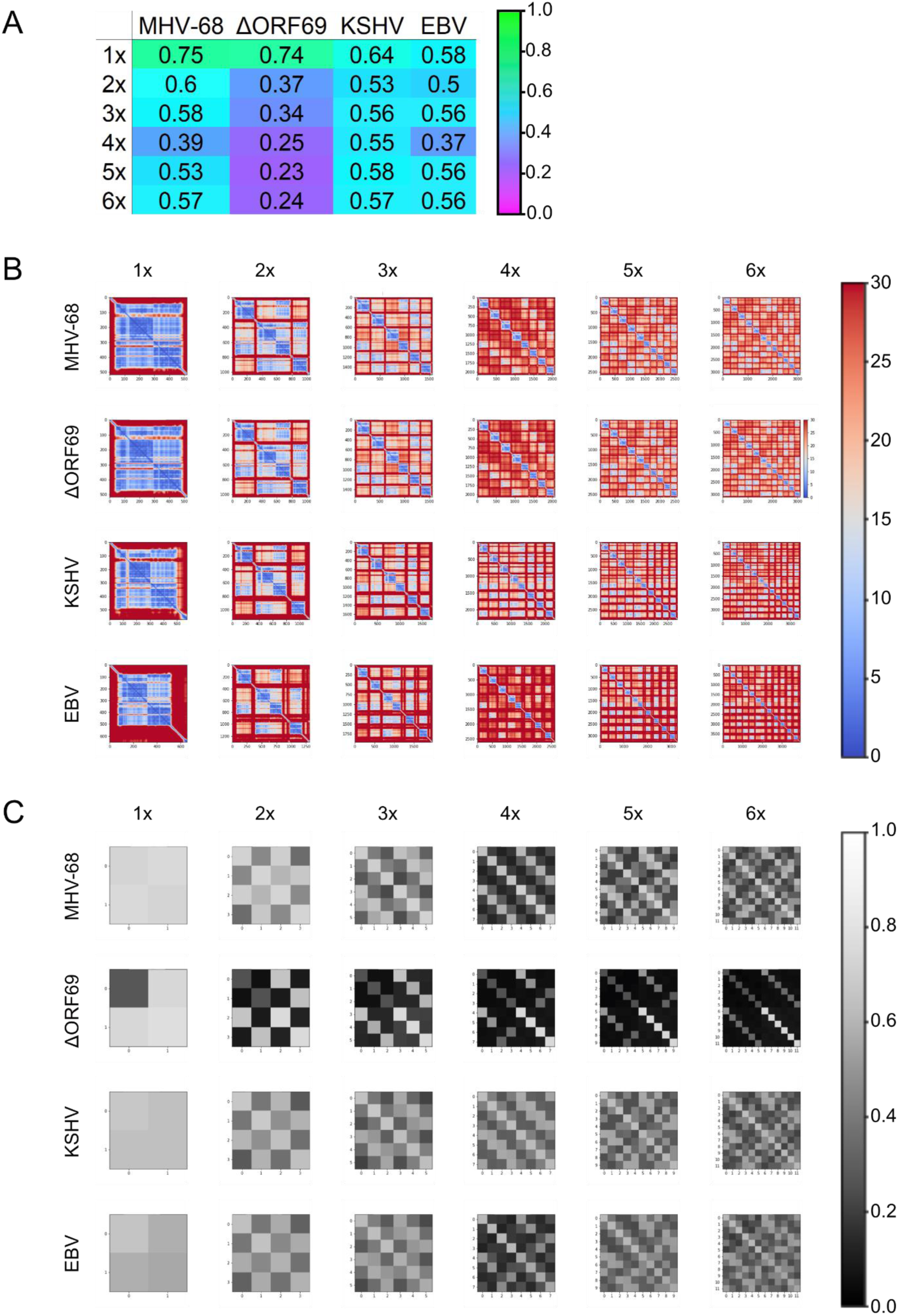
Quality assessment of AlphaFold 3 predictions of NEC oligomers of γ-herpesviruses. Different AlphaFold 3 quality scores for NEC-heterodimer oligomers (heterodimer to heterododecamer) in MHV-68, ΔORF69, KSHV, and EBV. (A) Global interface predicted Template Modeling (ipTM) scores for NEC-heterodimer oligomers (heterodimer to heterododecamer). The global ipTM score is color-coded from magenta (low confidence: 0) to green (high confidence: 1). (B) Predicted Aligned Error (PAE) plots for the NEC oligomers. The heatmap represents the expected distance error in Å between residue pairs, with colors ranging from red (high error: 30 Å) to blue (low error: 0 Å). (C) A pairwise comparison of ipTM scores between proteins is displayed as a grayscale heatmap of a 2D array. The scale ranges from black (low confidence: 0) to white (high confidence: 1).

In summary, we successfully designed a fluorescent ORF69 mutant of MHV-68, which retains its ability to interact with ORF67 but cannot oligomerize into higher-order complexes. Deleting the N-terminal region of ORF69 led to a frameshift mutation that preserved the first 68 amino acids, ensuring minimal disruption of the genetic context. Structural modeling using AlphaFold showed that while the ORF67-ΔORF69 complex remains stable as a heterodimer, it does not form higher-order oligomers, unlike the wild-type ORF67-ORF69 heterodimer.

### ΔORF69 grows on non-complementing cells

Previous work characterized an ORF69 knockout mutant by transfecting BAC-DNA into human 293 cells, providing insights into early replication events (34). However, since this approach does not assess full replication dynamics, whether the mutant could sustain infection beyond a single round of infection remained unclear. To further characterize our ΔORF69 mutant, we reconstituted the BAC-DNA in mouse NIH3T3 cells trans-complementing ORF69, generated a virus stock, and used it for subsequent experiments unless indicated otherwise. To rule out contamination with WT, we infected NIH3T3 cells, expecting no spread if ORF69 were strictly essential. Surprisingly, the ΔORF69 mutant produced plaques, albeit at a much slower rate and significantly smaller size than WT (Figure 2A, 2B). We used a custom-made image-processing pipeline to quantify plaques across 49 merged fields of view (FOV) for each condition (Figure 2C, 2D). The average ΔORF69 plaque size area was approximately 8-fold smaller than WT (Figure 2E), yet the mutant virus retained the capacity to spread without a functional NEC.

**Figure 2:**
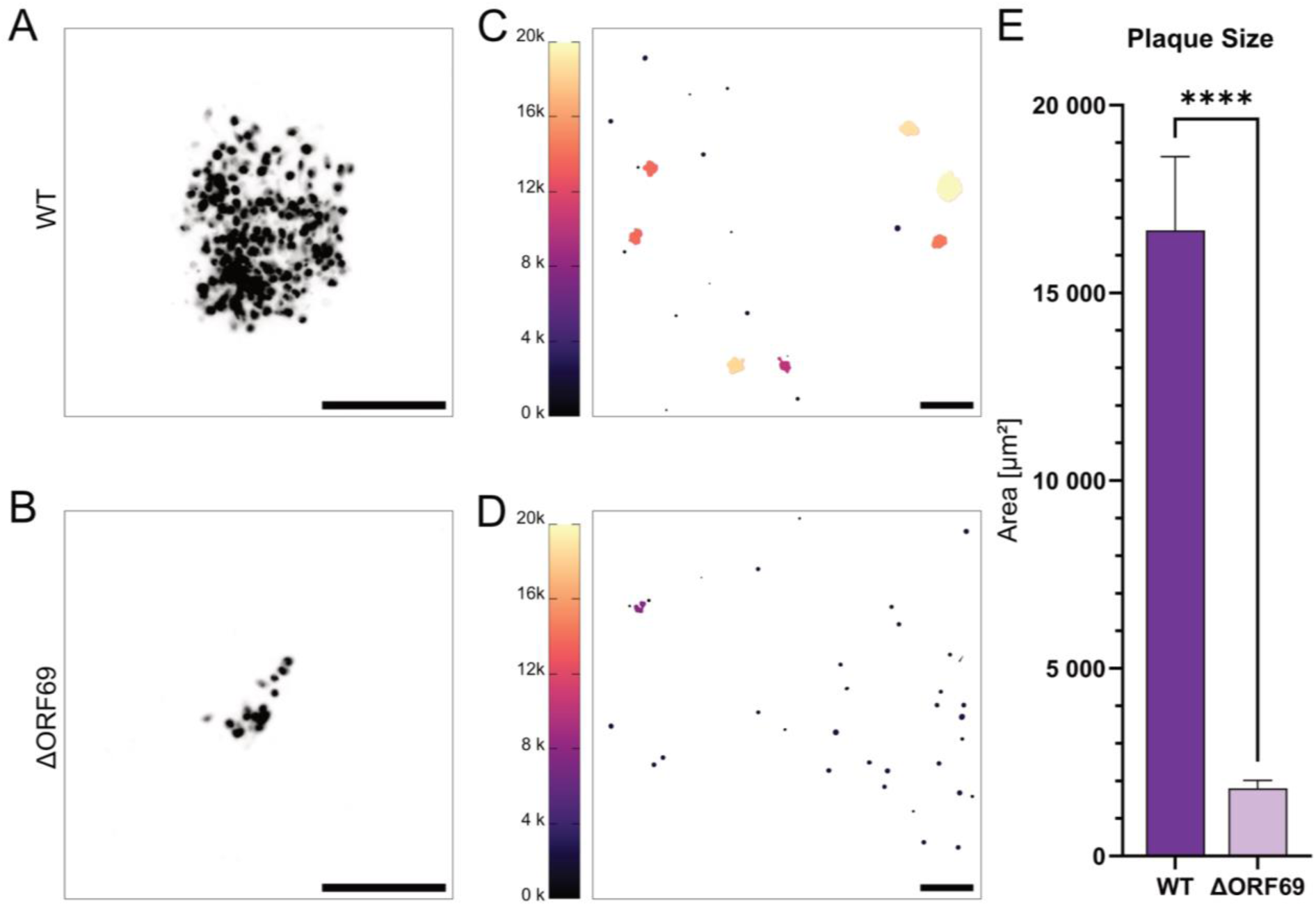
Comparison of WT and ΔORF69 plaque sizes. (A, B) Inverted fluorescent images of viral plaques 4 dpi for WT (A) and ΔORF69 (B); scale bar, 200 µm. (C, D) Binary mask outputs from the image-processing pipeline used to measure plaque area (µm²), color-coded according to plaque size, with the scale bar on the left. WT (C) and ΔORF69 (D); scale bar, 500 µm. Excluded values are color-coded in black. (E) Mean area distribution represented as bar graphs of three independent biological replicates, for WT (n=516) and ΔORF69 (n=182) plaques, with 95% confidence intervals shown. Kolmogorov-Smirnov test p-value: <0.0001.

We initially suspected a revertant because NEC proteins are thought to be essential for virus spread in most herpesviruses (39,40). However, Sanger sequencing of the genomic region of ORF69 and ORF67 of ΔORF69 showed no reversion or additional mutations in ORF69 or ORF67 (Figure S3A, S3B), indicating the mutant phenotype was maintained. These findings suggest that a fully functional NEC is not required for MHV-68 egress, as ΔORF69 exhibits drastically reduced plaque formation compared to WT but still retains replication and spread capacity, leading to further investigation into alternative mechanisms enabling capsid nuclear egress in the absence of a functional NEC.

**Figure S3:**
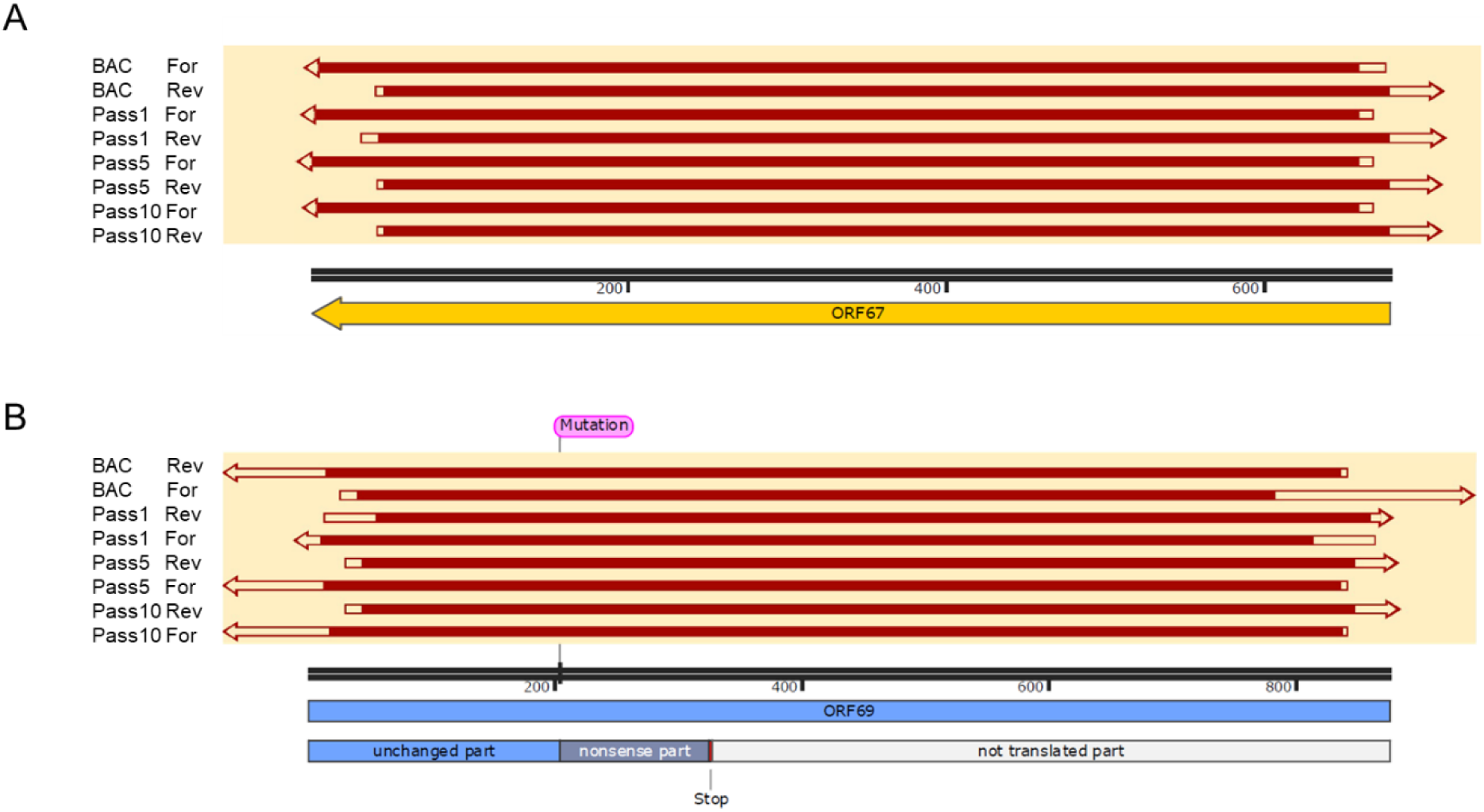
Alignment of Sanger sequencing results for ORF67 and ORF69. BAC, Pass1, Pass5, and Pass10 of ΔORF69 were sequenced. (A) Sequence alignment of ORF67 across the various virus passages. (B) Sequence alignment of ORF69 across the different virus passages, including WT ORF69 and the ΔORF69 deletion. Below each alignment, the corresponding ORF is shown: (A) ORF67 and (B) WT ORF69 and ΔORF69. Successful alignment with the reference sequence is indicated in red within the arrow.

### ΔORF69 does not acquire compensatory mutations

Previous studies on PrV ΔUL31 and PrV ΔUL34 knockout mutants, homologs of ORF69 and ORF67, respectively, showed that specific mutations accumulated when these PrV mutant viruses were passaged on non-complementing cells, leading to NEBD and increased titers (41,42). To investigate whether ΔORF69 also accumulates mutations like PrV that could bypass the NEC, we serially passaged ΔORF69 on non-complementing NIH3T3 cells. After ten passages (ΔORF69 Pass10), the cytopathic effect (CPE) was induced earlier, and the virus lost its fluorescence. To ensure the integrity of ORF67 and ORF69, we performed PCR amplification and Sanger sequencing on the viral DNA from the BAC, Pass1, Pass5, and Pass10 passages. Sequence alignment with ORF67 and ORF69 wild-type sequences confirmed the absence of mutations in these ORFs (Figure S3A).

Subsequent Illumina MiSeq platform-based sequencing and assembly with a reference-guided approach of the viral DNA of ΔORF69 Pass10 identified a 9 kb deletion at the 5’ end of the genome, including all eight tRNAs and the genes M1 to M4. No additional mutations were detected. To determine whether ORF4, located at the edge of the deletion, remained intact, we performed PCR amplification and subsequent Sanger sequencing. The successful amplification and sequence confirmation indicate that ORF4 is present in the P10 population (Figure 3, ΔORF69 Pass10). Large deletions at the 5’ end of the genome have been previously observed to occur spontaneously when MHV-68 is passaged on NIH3T3 cells (43). Notably, other field isolates, like MHV-72 and MHV-76 (Figure 3, MHV-72, MHV-76), also lack this region. MHV-72 lacks all tRNAs and the M1-M3 genes (44), while MHV-76 shares the same 5’ deletion profile (45,46) as ΔORF69 Pass10. In the parent virus of ΔORF69 and WT, the dispensable M1-region contains a duplicated ORF65, endogenously tagged with mCherry (35). Since this region is part of the 9 kb deletion observed during passaging in ΔORF69 Pass10, it explains the loss of fluorescence.

**Figure 3:**
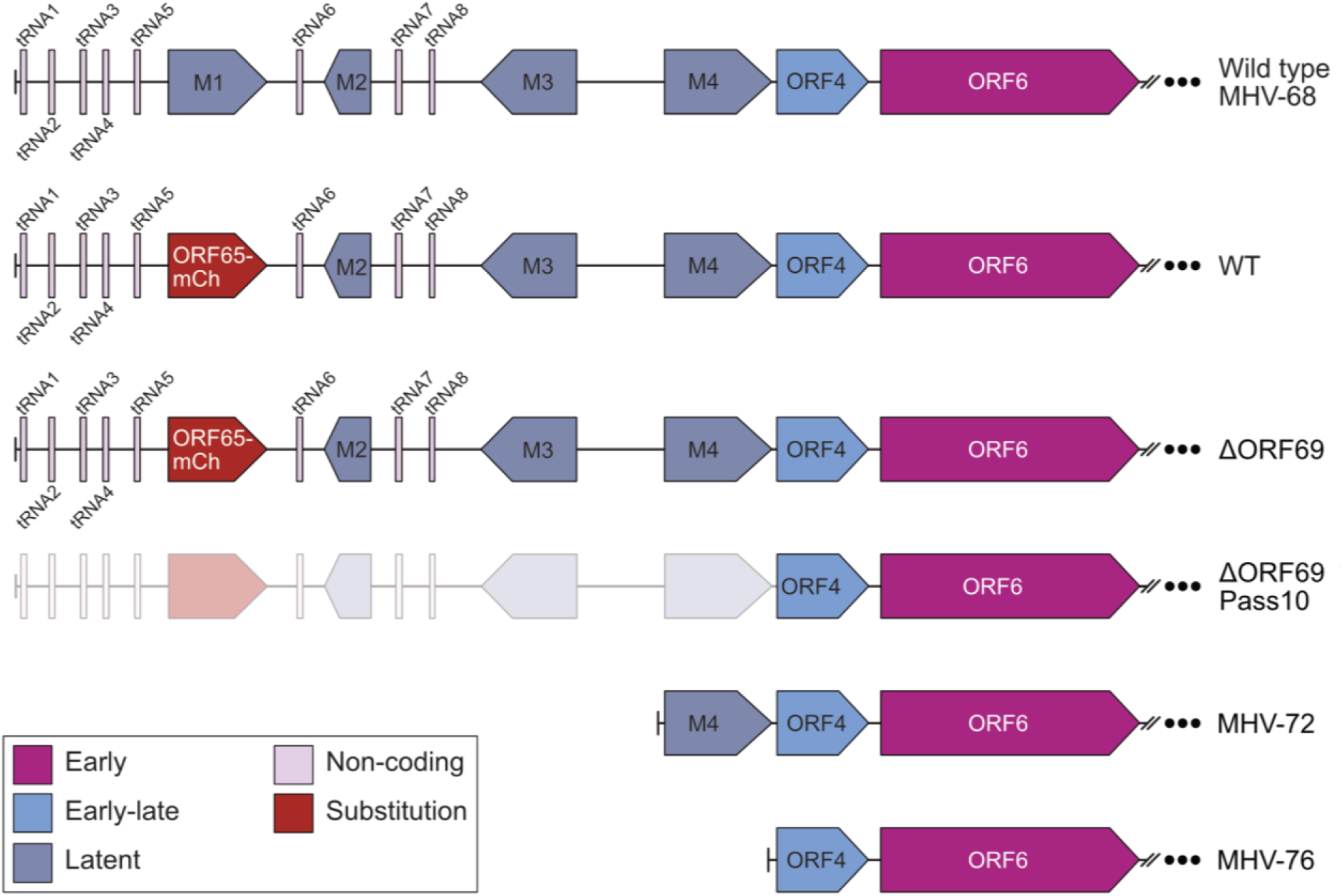
Genome comparison of ΔORF69 Pass10. The panel compares the genome of ΔORF69 Pass10, wild-type MHV-68, WT (this study), ΔORF69 (this study), and the field isolates MHV-72 and MHV-76. The ORF65-mCherry tag replacing the M1 gene is highlighted in red in the WT genome. The ΔORF69 Pass10 mutant exhibits a 9 kb deletion at the 5’ end of the genome, encompassing all eight tRNAs and the M1–M4 genes, shown as grayed out. This deletion is similar to those observed in field isolates MHV-72 and MHV-76. Non-coding tRNAs are depicted in lavender, latent genes in steel blue, early-late genes in cornflower blue, and early genes in deep magenta.

In contrast to PrV ΔUL31 and PrV ΔUL34, which acquired compensatory mutations upon serial passaging, ΔORF69 did not accumulate mutations that could facilitate alternative nuclear egress through NEBD. The absence of additional mutations, especially in the NEC-proteins or in glycoproteins, like observed for PrV (47), suggests that the observed increase in cytopathic effect is not due to the emergence of an intrinsically adapted subpopulation but instead reflects a general adaptation to passaging conditions. However, it remains possible that the mutant itself contains an inherent replication competence independent of adaptation. Given the lack of complementary mutations and the spontaneous deletion of dispensable regions, further passaging was unlikely to yield insights into alternative egress mechanisms.

### ΔORF69 capsids are restricted to the nucleus, but the virus does not show a packaging defect

With genetic compensation ruled out, we next investigated whether structural defects in viral assembly could contribute to the reduced spread of ΔORF69. The ORF69 homologs in related herpesviruses play a role in DNA packaging, with deletion of HSV-1 UL31 in α-herpesviruses leading to reduced overall viral DNA levels and impaired concatemeric DNA cleavage (48). In another mouse herpesvirus, β-herpesvirus MCMV, the homolog M53 may also play an accessory role in the DNA packaging of capsids (49). However, in the γ-herpesvirus EBV, deletion of BFLF2 does not alter monomeric genome levels but reduces the proportion of genome-filled C-capsids in the nucleus while increasing the abundance of abortive capsid forms, including A-capsids and B-capsids (50). To determine whether ΔORF69 shows a phenotype similar to HSV-1 or EBV, we performed transmission electron microscopy (TEM) in combination with serial sectioning on WT- and ΔORF69-infected cells to analyze at least 10 nuclei per condition across multiple depths, ensuring a robust quantification of nuclear capsid forms. We aimed to assess potential defects in capsid assembly or DNA packaging that might contribute to the observed reduction in viral spread.

The overall nuclear morphology was comparable between WT- and ΔORF69-infected cells. Both displayed regions of condensed chromatin adjacent to intact nuclei, with no observable lesions or disruptions in the ΔORF69 samples (Figure 4A, 4B). All three capsid forms were evenly distributed within the nucleoplasm in small, loose clusters, with no significant morphological differences between the two conditions (Figure 4C, 4D).

**Figure 4:**
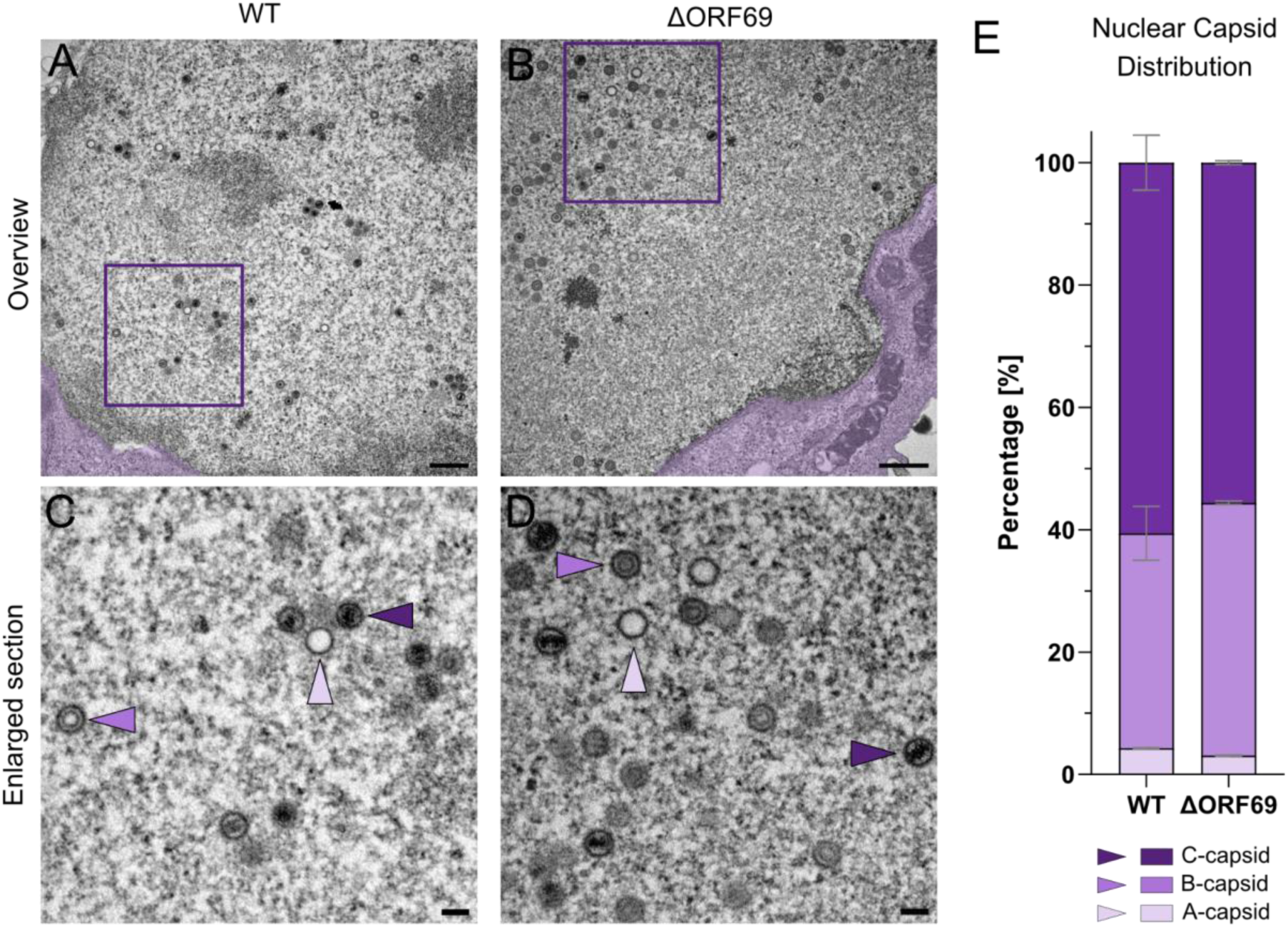
Quantification of nuclear capsid forms in WT and ΔORF69. (A, B) Representative TEM images of the nucleus in WT- and ΔORF69-infected NIH3T3 cells, respectively. The adjacent cytoplasm is false-colored in purple. Scale bar: 500 nm. (C, D) Enlarged views of the nucleoplasm from the boxed regions in (A) and (B) highlighting the three distinct capsid types: C-capsids (DNA-filled, indigo arrowheads), B-capsids (scaffold-containing, orchid arrowheads), and A-capsids (empty, lavender arrowheads). Scale bar: 100 nm. (E) Quantification of the different capsid forms. WT: Two independent biological replicates, n_cells_= 14, n_FOV_ = 455, n_capsids_= 5527, ΔORF69: Two independent biological replicates, n_cells_= 10, n_FOV_= 330, n_capsids_= 4761.

The quantification of nuclear capsid forms (Figure 4E) revealed a similar distribution between WT and ΔORF69. In both viruses, C-capsids, indicative of fully packaged, DNA-containing capsids, formed the majority (WT: 60.6% ± 4.5%; ΔORF69: 55.9% ± 0.3%). B-capsids, which represent abortive intermediates with retained scaffolds due to not initiated packaging, were slightly more abundant in ΔORF69-infected cells (WT: 35.1% ± 4.4%; ΔORF69: 41.3% ± 0.3%), while A-capsids, reflecting unsuccessful DNA packaging, comprised only a small fraction in both strains (WT: 4.3% ± 0.1%; ΔORF69: 3.1% ± 0.1%).

These findings suggest that the absence of ORF69 does not disrupt capsid assembly or DNA packaging, as shown by the comparable ratios and morphology of nuclear capsids in WT- and ΔORF69-infected cells. Moreover, we found no structural anomalies in ΔORF69-infected nuclei, leaving the mechanism by which capsids egress from the nucleus unresolved. Given that C-capsid abundance in the nucleus was unaffected, the reduced viral spread cannot be attributed to a defect in DNA packaging, as proposed for HSV-1 and EBV (48,50). To pinpoint the stage of viral replication affected by the ORF69 deletion, we next examined the distribution of capsids in the cytoplasm.

### Capsids are strongly reduced from the cytoplasm of ΔORF69 infected cells

To determine whether capsids can egress into the cytoplasm in very small quantities via a non-canonical route, potentially through NEBD as seen in PrV and HSV-1 (41,42,51,52), dilated nuclear pores, as implied for HSV-1 (26), or another unidentified mechanism, we examined the cytoplasm of WT- and ΔORF69-infected cells for the presence of capsids and their composition of capsid forms.

Unlike the nuclear morphology, the cytoplasmic morphology differed drastically between the two viruses. In WT-infected cells, the cytoplasm contained capsids, predominantly C-capsids (Figure 5A, indigo arrowhead), with only a small portion of A-capsids (Figure 5A, lavender arrowhead). At the same time, B-capsids were absent across all examined FOV. Dark, protein-rich structures were also observed, likely serving as tegument reservoirs. These structures appeared to play a role in the maturation process, as illustrated by three representative C-capsids in Figure 5A: a single C-capsid (bottom, indigo arrowhead), a C-capsid budding into a protein-rich structure (middle), and a fully tegumented and enveloped capsid, possibly following secondary envelopment (top). Quantification of these capsids showed a majority of C-capsids (96.6% ± 1.8%), with a minor fraction of A-capsids (3.3% ± 1.8%) (Figure 5E).

**Figure 5:**
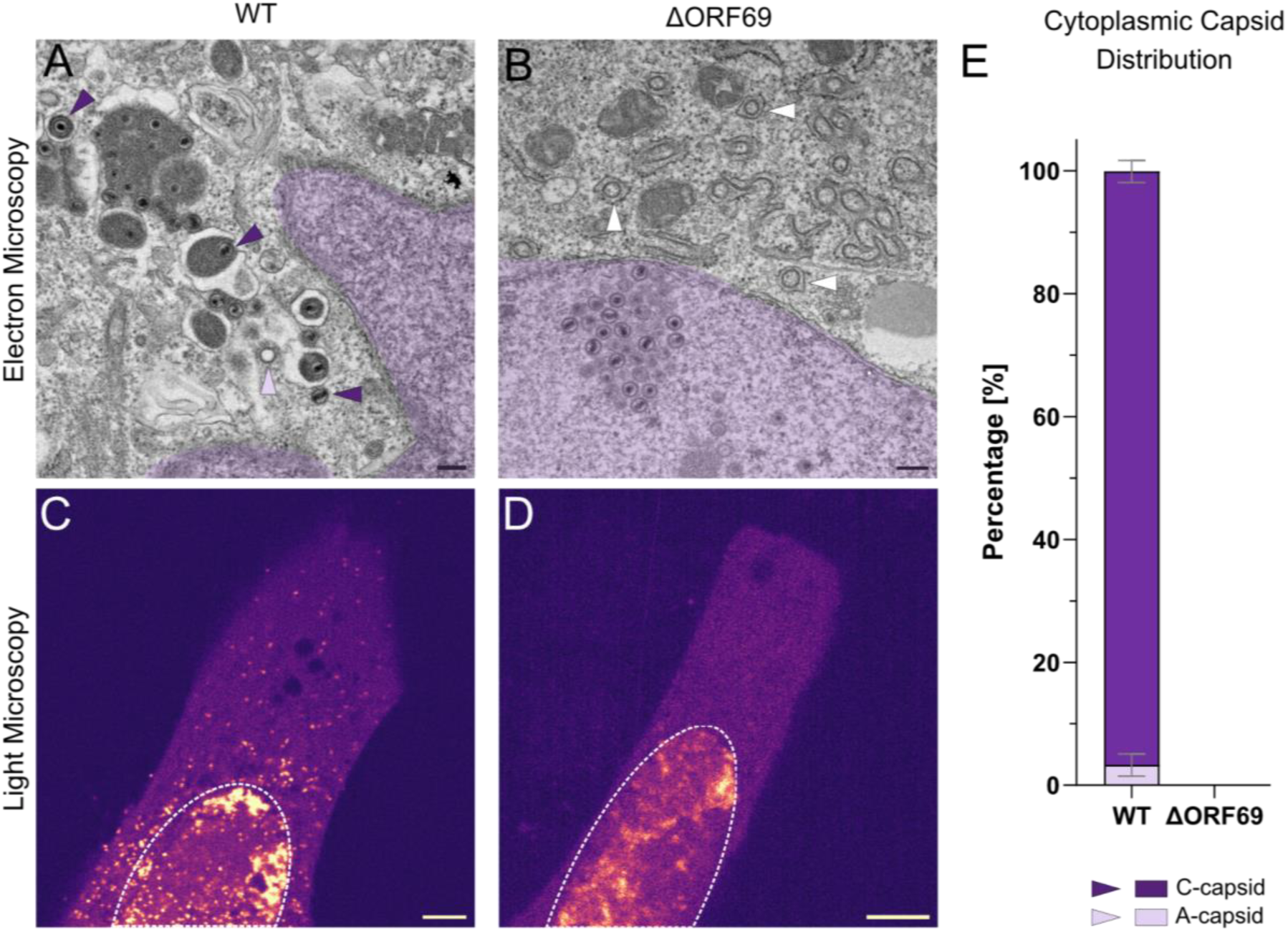
Analysis of cytoplasmatic capsids in WT and in ΔORF69. (A, B) Representative overview EM images of the cytoplasm of WT (A) and ΔORF69 (B) infected NIH3T3 cells. The adjacent nucleus is false-colored in purple. (A) C-capsids (DNA-filled, indigo arrowheads) and A-capsids (empty, lavender arrowhead). (B) Empty vesicular cytoplasmic structures (white arrowheads) Scale bar 200 nm. (C, D) Representative overview live-cell images of the NIH3T3 cells infected with WT (C) or ΔORF69 (D) at 1 dpi. The nucleus is marked with a dotted line. The images were color-coded with the “mpl-magma” LUT in Fiji ImageJ, where low-intensity signals are shown in indigo and high-intensity signals in yellow-white. Scale bar 5 µm. (E) Quantification of the different capsid forms in the cytoplasm of WT in EM. Two independent biological replicates, n_cells_= 14, FOV: 219, n_capsids_= 1956.

In contrast, no cytoplasmic capsids were detected in ΔORF69-infected cells across all examined fields of view. Instead, the cytoplasm contained enveloped, empty vesicular structures (Figure 5B, white arrowhead) absent in WT cells. In some fields of view where parts of the nucleus were visible, small clusters of C-capsids were observed near the nuclear envelope. However, no direct contact was detected between capsids and the nuclear envelope, which would indicate interaction with it. Interestingly, these vesicular structures resemble those observed in cells expressing KSHV ORF67 and ORF69 in SF21 cells (53). Desai *et al.* (2012) reported that co-expression of ORF67 and ORF69 resulted in numerous virion-sized circular vesicles, a phenotype also described for PrV homologs (54). The similarity in vesicle formation suggests that the functional N-terminal region of ORF69 may be sufficient to induce these structures since the expression of only ORF69 in KSHV does not induce these cytoplasmic structures (53).

We performed live-cell imaging to investigate whether capsids were present in the cytoplasm at earlier time points than those used for EM analysis, allowing a higher throughput of cells than EM. At 1 dpi, numerous fluorescent capsids were observed in the cytoplasm of WT-infected cells (Figure 5C). However, no fluorescent capsids were detected in the cytoplasm of ΔORF69-infected cells, and only the signal in the nucleus was visible (Figure 5D). This absence of cytoplasmic capsids in ΔORF69-infected cells at both 1 dpi in live-cell imaging and 4 dpi in EM supports the hypothesis that capsid egress in ΔORF69-infected cells occurs very rarely, making it challenging to capture the exact time point of nuclear egress.

To determine whether capsids were present only in the cytoplasm at a specific stage of infection, we infected a monolayer of cells with ΔORF69 for 21 days until visible plaques appeared. Since we aimed to capture all phases of the infection cycle, we specifically processed plaques for TEM, reasoning that this approach would maximize the likelihood of observing various viral life cycle stages. However, the strong tendency of ΔORF69-infected cells to round up and detach poses a technical challenge for capturing late-stage infection events. We used enhanced crosslinking by prolonged glutaraldehyde exposure after the standard fixation period before further processing for TEM to stabilize the cells as much as possible. While most infected cells had intact nuclei (Figure S4A, S4B), we successfully captured infected cells in the round-up state, characterized by severely compromised nuclei and highly altered overall cell morphology (Figure S4C). Additionally, we identified a cluster of cytoplasmic capsids with all capsid forms present (Figure S4D). However, it was not possible to quantify and draw reliable conclusions beyond the presence of a few cytoplasmic capsids of different capsid forms due to the small number of cells in this particular state and the few reliably identifiable cytoplasmic capsids.

**Figure S4:**
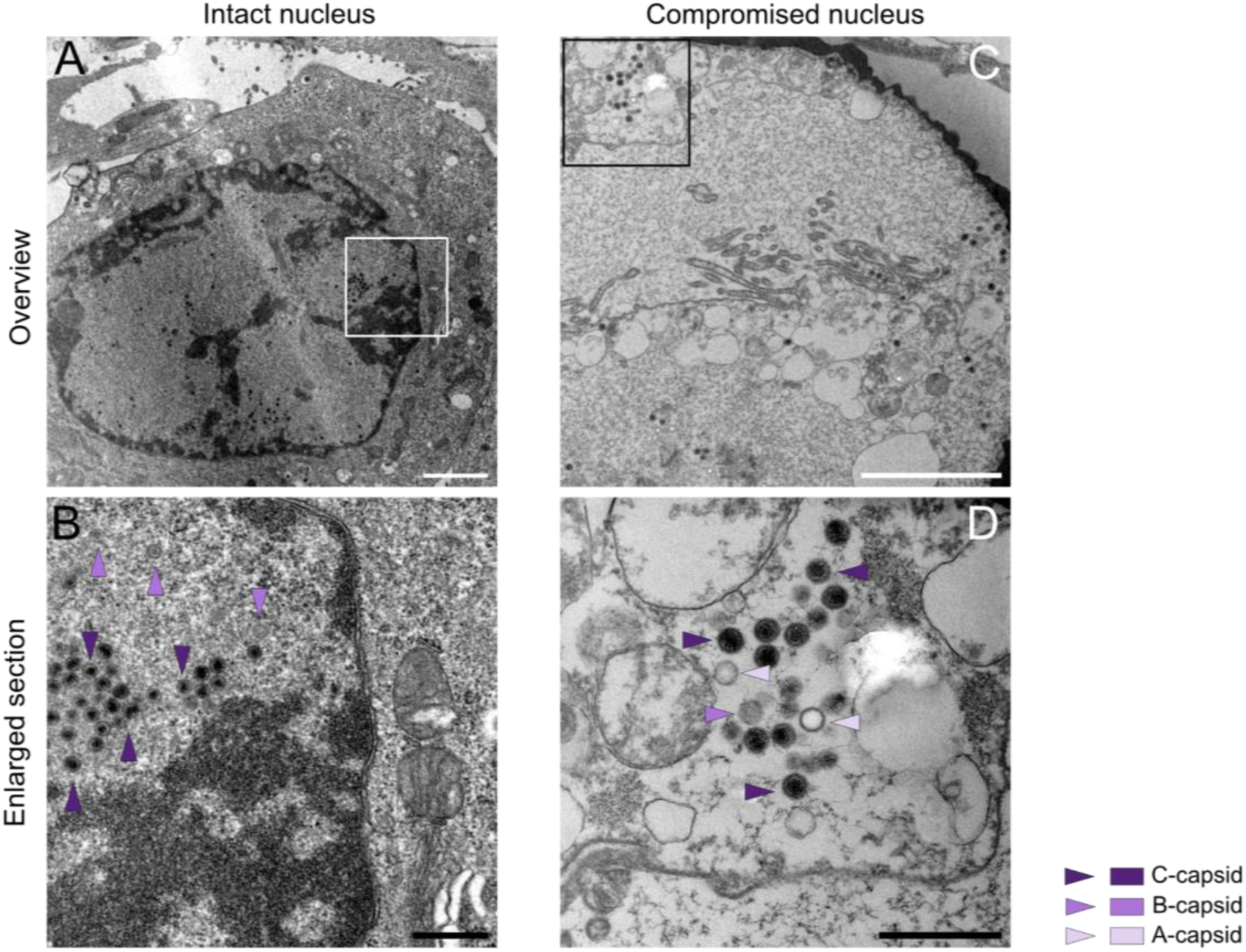
Late-stage cytoplasmic capsids in ΔORF69. (A) Representative EM overview of a ΔORF69-infected NIH3T3 cell at 21 dpi, showing an intact nucleus. (B) Enlarged section of (A) (marked with a white square), highlighting retained nuclear capsids, primarily C-capsids (indigo arrowheads) and B-capsids (orchid arrowheads). (C) Overview of a ΔORF69-infected cell with a compromised nucleus. (D) Enlarged section of (C) (marked with a black square), showing a cytoplasmic capsid cluster containing all capsid forms: DNA-filled C-capsids (indigo arrowheads), scaffold-containing B-capsids (orchid arrowheads), and empty A-capsids (lavender arrowheads).

These observations suggest that in MHV-68, nuclear egress of C-capsids is tightly regulated by the NEC, acting as a quality control checkpoint that selectively permits only mature, DNA-containing C-capsids to egress, as seen in WT-infected cells (Figure 6A, 6E). This selective mechanism underscores the crucial role of the γ-herpesvirus NEC in the nuclear egress of mature capsids. In contrast, in ΔORF69-infected cells, the translocation of capsids into the cytoplasm still occurs, but only in cells with compromised nuclei. Given that this process likely follows an unknown pathway in these severely compromised cells, it remains uncertain whether secondary envelopment, an essential cytoplasmic step in the herpesvirus life cycle, can proceed efficiently. It is not clear whether the severely compromised state of ΔORF69-infected cells allows them to support regular cellular functions. Consequently, viral processes that exploit host factors, such as secondary envelopment (55,56), may be impaired. It remains to be determined whether ΔORF69-infected cells can produce fully enveloped C-capsids.

**Figure 6:**
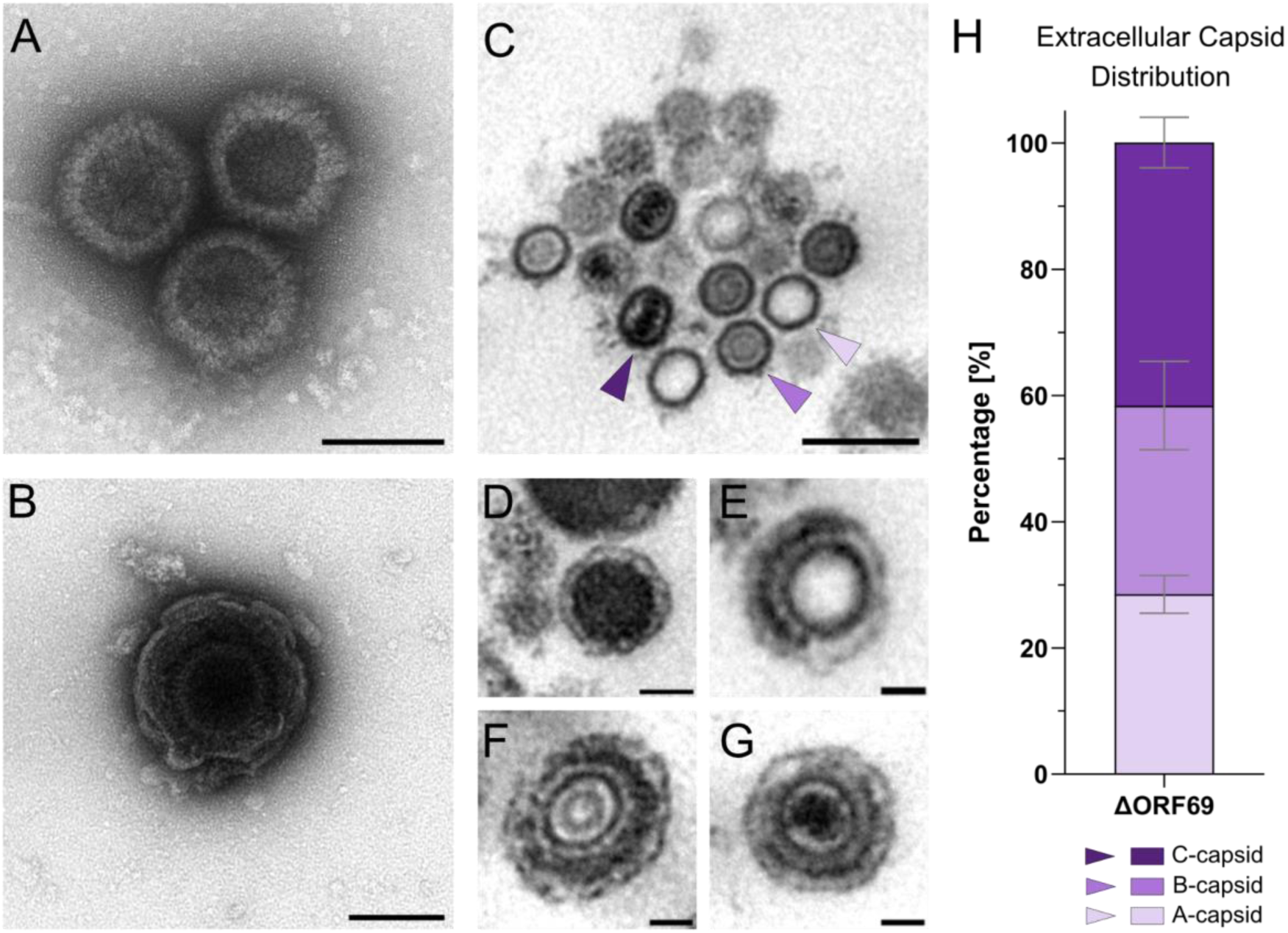
Extracellular enveloped capsid forms of ΔORF69. (A, B) Negative staining of pelleted virus supernatant for quality control, showing intact particles. (A) A cluster of three intact capsids. (B) A single enveloped capsid. (C-G) Overview EM images of particles from the supernatant, embedded in epoxy resin and prepared for ultra-thin sectioning. (C) A cluster of naked capsids displaying all three capsid forms: DNA-filled C-capsids (indigo arrowhead), scaffold-containing B-capsids (orchid arrowhead), and empty A-capsids (lavender arrowhead). (D) Dense body. (E) Enveloped A-capsid with an incomplete tegument layer. (F) Enveloped B-capsid. (G) Enveloped C-capsid. (H) Quantification of the different capsid types present in the ΔORF69 supernatant. Two independent experiments, n_Grids_= 3, n_FOV_= 178, n_capsids_= 1550.

### ΔORF69 produces a small fraction of infectious particles

Herpesvirus capsids require envelopment to become infectious. Since we observed few cytoplasmic particles in ΔORF69-infected cells after nuclear membrane disintegration, with no enveloped particles present, we sought to determine whether the supernatant from these cells contained potentially infectious, enveloped C-capsids. Given the technical challenges we encountered with processing plaques for TEM, which resulted in a limited number of cells for analysis, we focused on the supernatant to better assess the presence of infectious particles.

We pelleted the virus from the supernatant of ΔORF69-infected cells and performed negative staining as a quality control step (Figure 6A, 6B). The pellet contained unenveloped capsids (Figure 6A) and some enveloped capsids (Figure 6B). Based on the consistency of the pellet, we either processed the pellet directly by ultra-thin sectioning or utilized cellulose capillary tubes for support prior to sectioning. Capsids often appeared in clusters or small aggregates of different capsid forms (Figure 6C). Besides capsids, the majority of particles we observed were various sizes of electron-dense vesicles, likely representing dense bodies. These non-infectious particles consist primarily of tegument protein and do not contain a capsid (Figure 6D). Surprisingly, we detected all three capsid forms enveloped (A-capsid Figure 6E, B-capsid Figure 6F, and C-capsid Figure 6G), albeit at low quantities. Some enveloped particles only showed a partial tegument layer between the capsid and envelope (Figure 6E). Quantification of the individual capsid forms indicated a balanced distribution of capsid types in the supernatant, with C-capsids forming the majority (41.7% ± 4.0%), followed by B-capsids (29.9% ± 7.0%) and a high fraction of A-capsids (28.5% ± 3.0%) (Figure 6H).

These results suggest that capsids can egress from the nucleus without a functional NEC after the nuclear membrane disintegrates. Likely, some capsids can acquire a tegument layer and an envelope after the nuclear membrane is compromised, regardless of the capsid form. The presence of the different enveloped capsid forms suggests that the process of secondary envelopment can proceed without regard for a specific capsid form or content and, therefore, possesses no intrinsic quality control. In contrast, the NEC acts as a stringent quality control checkpoint, restricting nuclear egress almost exclusively to mature, DNA-containing C-capsids. While enveloped A- and B-capsids are non-infectious due to the absence of viral genomes, they retain the ability to fuse with host cells. However, these particles cannot initiate transcription without the genome, which is crucial for suppressing antiviral immune responses. Consequently, if these non-infectious particles fail to counteract host defenses, the immune response triggered may reduce the overall effectiveness of the remaining small fraction of infectious C-capsids, thereby limiting viral spread even more.

### Block of CDK-dependent pathway decreases WT, and ΔORF69 spread

Given that the precise mechanism of nuclear membrane disintegration in ΔORF69 remains unsolved, we investigated whether pseudo-mitotic processes might facilitate an alternative route for nuclear egress. For this, we treated WT- and ΔORF69-infected cells with either Roscovitine, a cyclin-dependent kinase inhibitor, or U0126, a MEK1/2 inhibitor. This strategy is based on previous work on PrV ΔUL31Pass and ΔUL34Pass mutants, which regained near wild-type replication through serial passaging and induction of NEBD (41,42). The residual infectivity in PrVΔUL31 or PrVΔUL34 enabled serial passaging, suggesting that the virus possesses an intrinsic capacity for low-level spread. Although no compensatory mutations were detected in our ΔORF69 mutant after passaging, these findings raise the possibility that an alternative egress mechanism may be conserved among herpesviruses that can be exploited under certain conditions. Roscovitine potently inhibits multiple CDK complexes with IC₅ ₀ values of 0.65 μM for cdc2/cyclin B, 0.7 μM for cdk2/cyclin A, 0.7 μM for cdk2/cyclin E, and 0.16 μM for cdk5/p53. At the same time, its inhibition of ERK1/2 is considerably weaker, with IC₅ ₀ values of 34 μM and 14 μM, respectively (57). Roscovitine has been shown to reduce viral replication in wild-type MHV-68 (58). In contrast, U0126, with IC₅ ₀ values of 0.07 μM for MEK1 and 0.06 μM for MEK2 (9), selectively targets the MAPK/ERK pathway and has been shown to reduce viral replication in wild-type MHV-68 only slightly (59). Prior to the experiments, we assessed cytotoxicity (Figure S5) and selected concentrations that maximized inhibition of cell proliferation, leading to a stable cell number throughout the testing period. The stable cell numbers were based on the mitotic arrest induced by Roscovitine in the G0, G1, S, and G2/M phases and the delay in early/mid-G2 progression induced by U0126. Consistent with previous studies, these concentrations effectively modulated distinct cell cycle checkpoints while preserving overall cell viability. Cells were treated daily with 5 μM Roscovitine, 50 μM U0126, PBS, or vehicle control DMSO.

**Figure S5:**
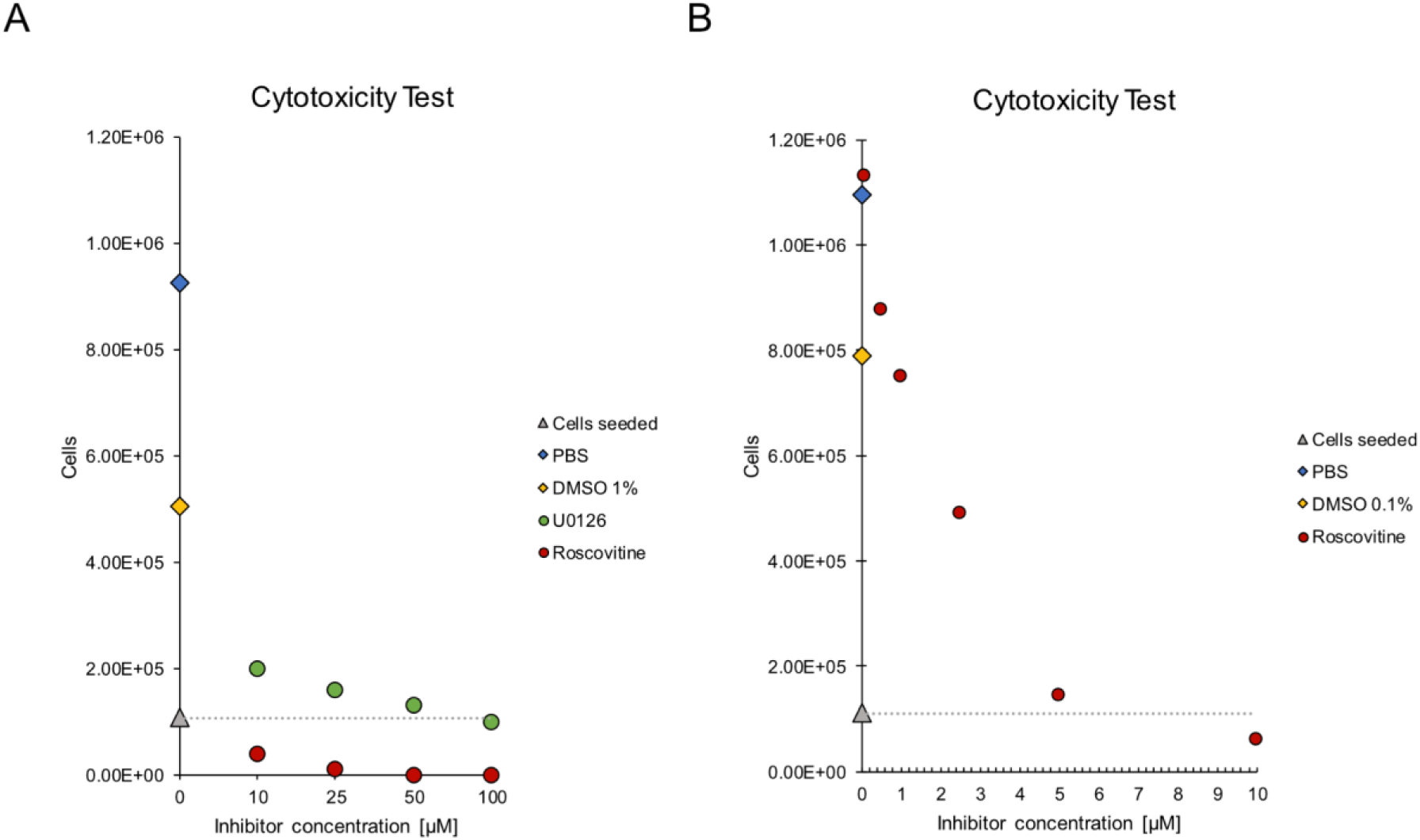
Cytotoxicity analysis of U0126 and Roscovitine on NIH3T3 cells. NIH3T3 cells were treated with varying concentrations of U0126 and Roscovitine, and cell viability was assessed at 3 dpi. (A) High concentrations of Roscovitine (10– 100 µM) and all U0126 concentrations (10–100 µM). (B) Low concentrations of Roscovitine (0.1–10 µM). The gray dotted line represents the initial seeding density. Blue indicates the PBS control, yellow represents the corresponding DMSO control, green (Panel A only) shows U0126-treated cells, and red shows Roscovitine-treated cells. The y-axis displays the final cell numbers at 3 dpi, while the x-axis represents inhibitor concentrations. All controls are aligned at 0 µM.

Fluorescent plaques were quantified using the image processing pipeline described earlier (Figure 7A). In WT infections, U0126 and Roscovitine significantly reduced plaque size compared to controls (Figure 7B). The reduction observed with U0126 suggested that MEK1/2 signaling contributes to the efficient viral spread. In contrast, ΔORF69 plaque size was not affected by U0126 (Figure 7C), suggesting that ORF69 and MEK1/2 signaling function within the same pathway. The lack of an additional effect implies that the role of MEK1/2 in viral propagation may depend on ORF69. However, Roscovitine shows a significant inhibitory effect on both WT and ΔORF69 plaque formation. Although the effect of Roscovitine was more pronounced in WT compared to the respective control, its significant reduction of the already very small ΔORF69 plaques suggests that the CDK-dependent pathway is critical for residual viral propagation and likely acts through a distinct mechanism from MEK1/2.

**Figure 7:**
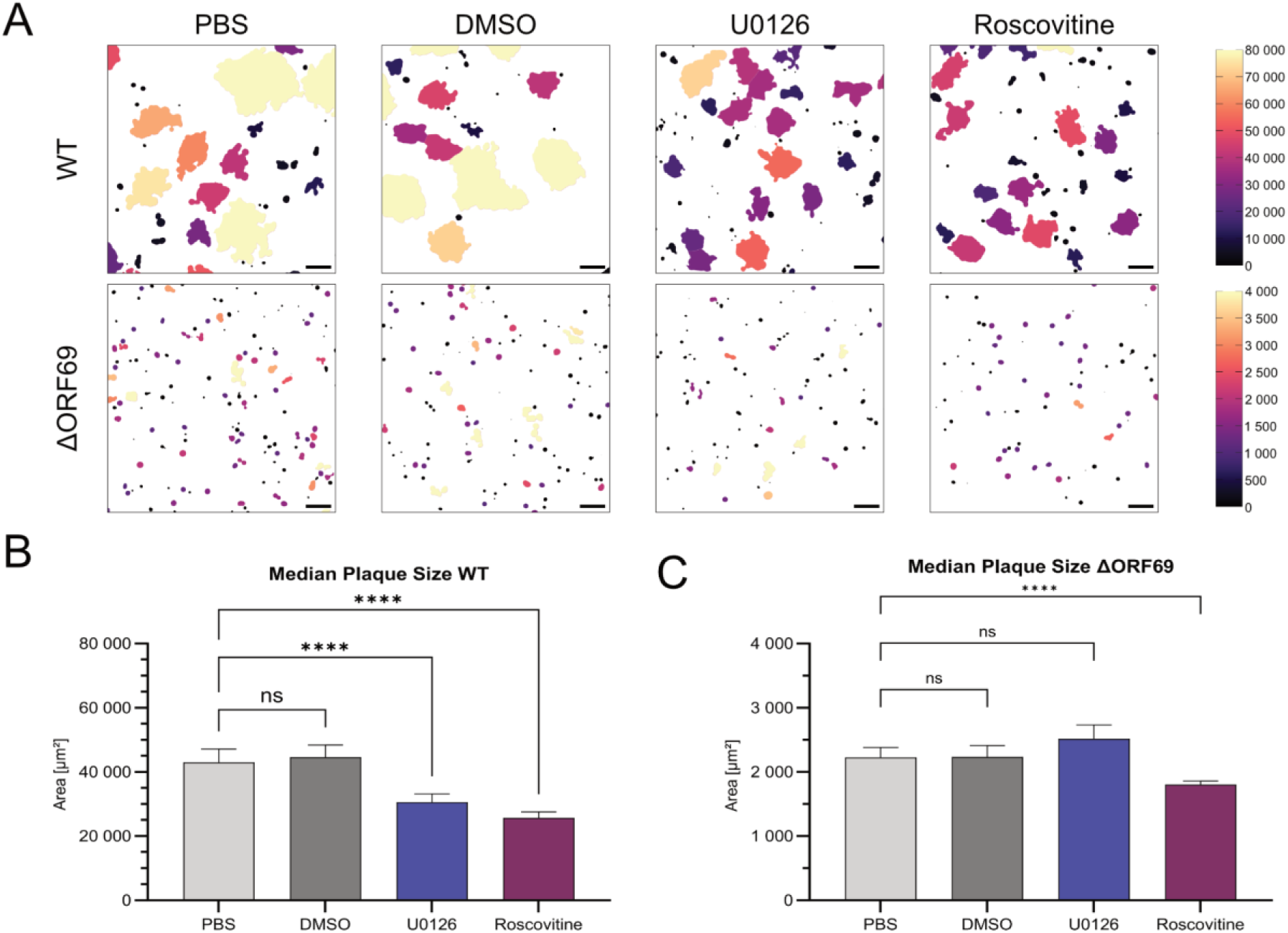
Effect of inhibitors on plaque size in WT and ΔORF69. (A) Binary mask outputs from the image-processing pipeline showing plaque area (µm²) for WT and ΔORF69 under different treatments. The upper panel displays WT plaques under PBS, DMSO, U0126, and Roscovitine, and the lower panel shows ΔORF69 plaques under the same conditions. The upper panel is color-coded according to the scale (0-80000 µm²), while the lower panel uses a 0-4000 µm² scale. Scale bar 200 µm. (B) Quantified plaque area (µm²) for WT plaques in four independent experiments (PBS_WT_ n=3817, DMSO_WT_ n=3274, U0126_WT_ n=2157, Roscovitine_WT_ n=2041). A threshold of >3000 µm² was applied. Kolmogorov-Smirnov test p-value: <0.0001. (C) Quantification of plaque area (µm²) for ΔORF69 plaques in four independent experiments (PBS_ΔORF69_ n=1130, DMSO_ΔORF69_ n=1269, U0126_ΔORF69_ n=1247, Roscovitine_ΔORF69_ n=1318). A threshold of >1000 µm² was applied. Kolmogorov-Smirnov test p-value: <0.0001.

## Discussion

Our study demonstrates that the NEC of murine gammaherpesvirus 68 in infection, specifically the nucleoplasmic protein ORF69, is not essential for nuclear egress but plays a crucial role in mediating C-capsid specificity. ΔORF69 retained the ability to replicate and spread in NIH3T3 cells, although with reduced efficiency and smaller plaque sizes than the parent virus. Quantitative electron microscopy showed that the absence of ORF69 led to a loss of selective nuclear egress, resulting in the presence of A-, B-, and C-capsids in the cytoplasm, which contrasts with parent virus infection, where only DNA-filled C-capsids are efficiently exported. The predominant export of C-capsids suggests that the γ-herpesvirus NEC functions as a quality control checkpoint, selectively facilitating the translocation of genome-filled C-capsids while retaining immature or defective capsids within the nucleus.

The role of the NEC as a membrane remodeler (53,54,60) and a translocator of capsids through the nuclear membrane in an envelopment-development manner is well established (61). In α-herpesviruses, the deletion of NEC components results in a complete block of nuclear egress and the accumulation of capsids within the nucleus (48,62–64). Similar phenotypes have been reported for the γ-herpesviruses EBV, where knockout of either BFLF2 or BFRF1 leads to nuclear retention of capsids (50). However, evidence suggests that the NEC is not essential, as residual infectivity has been detected in multiple herpesviruses lacking NEC components (31,41,42,52,65–68). Our findings in MHV-68, which align with previous transfection-based studies on NEC-knockout mutants (31), indicate that nuclear egress of capsids can proceed in the absence of a functional NEC, albeit in a less efficient and non-selective manner, as shown by the increased levels of extracellular A- and B-capsids in our study. The NEC has been discussed as a potential quality control mechanism for capsid selection, involving the relative abundance of the HSV-1 UL25, ORF19 in MHV-68 and KSHV, on the capsid. The UL25 copy numbers are the highest on C-capsids, followed by A- and B-capsids (3,69). The model proposes that the NEC might favor capsids with a high copy number of UL25 present, offering a molecular foundation for quality control. However, this model requires a highly accurate quality mechanism and could not be proven. An alternative hypothesis proposes that the asymmetric unit harboring the portal may be a key determinant for capsid selection (70). The portal in C-Capsids may have additional characteristics, such as the portal cap, an alternative structural arrangement of the portal itself, or a feature linked to the viral genome. One of these potential features could provide a structural element on the capsid preferentially selected by the NEC. However, we could not identify the underlying selection mechanism with this study design.

Furthermore, our observations highlight a critical distinction between nuclear and cytoplasmic quality control mechanisms. The presence of enveloped A-, B-, and C-capsids in the extracellular space suggests that secondary envelopment does not impose a strict quality control step concerning the capsid form. This implies that the NEC serves as the final quality checkpoint for genome-filled capsid selection in the herpesvirus life cycle.

Other proposed functions for ORF69 homologs in HSV-1 and PrV are that UL31 may guide capsids to the nuclear envelope and facilitate interactions with UL34 (71–76). This observation fits our observation that nuclear C-capsids cluster in the vicinity of the nuclear envelope but do not interact with it. So, while the residual N-term of ORF69 can interact with ORF67, it cannot interact with the capsid. Notably, the N-term of MHV-68 ORF69 and its homologs are highly variable, and different functions have been proposed (77). In HSV-2 UL31, this region is associated with effective DNA replication, packaging, and DNA damage response (36). Similarly, HSV-1 plays a role in membrane budding, suggesting that the N-terminal of ORF69 may also have these functions (37). Since we did not observe a packaging defect comparable to EBV (50), our findings suggest that the residual N-term region of ORF69 may play a role in packaging. However, further systematic investigation is required to confirm this.

Despite the lack of a functional NEC and quality control, ΔORF69 was still capable of viral spread, suggesting the existence of alternative nuclear egress mechanisms. The existence of alternative escape mechanisms raises questions about their evolutionary significance. If alternative nuclear egress were a purely ancestral mechanism, it would likely have been lost in modern herpesviruses due to selective pressure favoring efficient replication. Others have shown that allowing the capsids to enter the cytoplasm does not restore full replication capacity. Therefore, it’s likely that the NEC presents another maturation step in the life cycle (52). Instead, its retention as a potential backup strategy suggests an adaptive advantage, particularly in cell types or conditions where the NEC function is impaired. This is consistent with the idea that herpesviruses maintain redundant pathways to ensure survival under diverse conditions. While NEC-mediated nuclear egress remains the dominant mechanism, alternative routes may be utilized in specific contexts. There is evidence that there are underlying cellular mechanisms capable of exporting large cargo from the nucleus, and it has been hypothesized that herpesviruses may have hijacked this pathway (78). Another possible explanation for residual infection is the nuclear envelope disassembly. In cells, this process occurs during mitosis, but a serially passaged mutant of PrV was also able to induce this NEBD. However, this process was sensitive to inhibition of the CDK- or MEK1/2 pathways. Interestingly, the CDK inhibitor Roscovitine reduced the titers of the passaged viral mutants (41,42). The role of CDKs in herpesvirus replication is increasingly recognized, with several studies implicating CDK activity in viral gene expression and egress (79,80). Our findings that CDK inhibition also reduces viral spread in WT and ΔORF69 suggest that CDKs play a role beyond NEC function, possibly regulating additional steps in the viral life cycle. Moreover, herpesvirus-encoded cyclins, such as MHV-68 v-cyclin, have been shown to associate with CDK2 and CDC2, further underscoring the importance of host kinase regulation in viral replication (81). On the other hand, the MEK1/2 inhibitor U0126 reduced plaque size in our WT mutant without altogether abolishing plaque formation, suggesting that while the virus may modify MEK1/2 signaling to enhance efficient viral spread, as observed for PrV (82), it is not absolutely essential for plaque formation. In contrast, the ΔORF69 plaque size was not affected by U0126, indicating that MEK1/2 signaling does not play a significant role in viral spread when the NEC is non-functional.

Our findings contribute to growing evidence that herpesviruses employ NEC-dependent and alternative nuclear egress routes, with the NEC acting as a critical quality control checkpoint for capsid selection. The observation that secondary envelopment does not impose strict quality control further supports the notion that nuclear egress represents the last major checkpoint in the herpesvirus replication cycle. Although an alternative nuclear egress route appears to be a viable but highly inefficient and barely detectable option under certain conditions, the functional NEC remains the predominant mechanism for efficient and selective nuclear egress. The reduced replication efficiency and lack of compensatory adaptation in ΔORF69 emphasize the importance of NEC-mediated capsid selection in optimizing viral fitness. These insights enhance our understanding of herpesvirus nuclear egress mechanisms.

## Materials and Methods

### Structure prediction

For the structure prediction of the NEC heterodimers and heterooligomers of MHV-68, EBV, and KSHV, the protein sequences were downloaded from https://www.uniprot.org using the entry number found in the table of our HerpesFolds database (23). AlphaFold 3 (38) predictions were performed through the AlphaFold Server at https://alphafoldserver.com. UCSF ChimeraX version 1.8 (https://www.cgl.ucsf.edu/chimerax) was used to visualize structures. Data analysis was carried out with custom-written Python scripts, which were partly generated with the assistance of large language models and thoroughly tested manually, as previously published (23).

### Viruses

The virus mutants used in this study are based on the BAC backbone previously described (83,84). MHV-68 ORF65-mCherry, referred to as WT throughout this study, was generated and characterized as described previously (85). To insert the expression cassettes encoded by the above-described rescue plasmids into the MHV-68 BAC, we used the single-step FRT/Flp system originally established for MCMV (8). For this purpose, Escherichia coli strain DH10B (Invitrogen), containing the MHV-68 BAC-backbone and the temperature-sensitive Flp recombinase expression plasmid pCP20 (86), was transformed with various R6K-gamma-driven pOTO constructs (8) carrying ORF65-fusion cassettes. Transformation and selection were performed as previously described (87). The ΔORF69 BAC was generated using a two-step replacement procedure, as described (88), with the shuttle plasmid outlined above. Correct recombination was verified by analysis of the restriction patterns of the respective ΔORF69 and the WT-BAC by restriction digest with EcoRI, HindIII, and SacI for 1h at 37°C and gel electrophoresis on a 0.8% agarose gel run at 180 V for 2h and subsequently 30 V overnight.

### Cells

The experiments were carried out on the NIH3T3 mouse fibroblast cell line. For trans-complementing ORF69, NIH3T3 cells were transduced using a third-generation lentiviral system. To generate a lentiviral vector using the Gateway Recombination Cloning Technology, ORF69 was amplified, with flanking attB1 and attB2 sites. It was cloned into pDONR221 using a BP reaction according to the manufacturer’s protocol (Thermo Fisher, USA), yielding the entry clone. An LR reaction was then performed between the entry clone and the destination vector pLenti CMV Puro Dest (pLenti CMV Puro DEST (w118-1) was a gift from Eric Campeau & Paul Kaufman) (Addgene plasmid #17452) to generate the pLenti_CMVPuro_ORF69 construct. Lentiviral particles were produced by transfecting human 293XT cells with pLenti_CMVPuro_ORF69 and the necessary packaging plasmids. Virus-containing supernatants were harvested and used to transduce NIH3T3 cells. Transduced cells were subjected to puromycin selection to enrich for stable ORF69 expression. These cells were subsequently used to produce virus stocks of ΔORF69 under non-selective conditions. attB1_ORF69 F: 5’-GGGGACAAGTTTGTACAAAAAAGCAGGCTTAatgcgctcaacaggctctg-3’ attB2_ORF69 R: 5’-GGGGACCACTTTGTACAAGAAAGCTGGGTTttgctgagaaagacgagatacaatgttga-3’

### Plaque Size Measurement

3T3 cells were seeded on 6-well plates at a density of 5.4 × 10^5^ cells per well to achieve 80% confluency the following day. Cells were infected with serial dilutions (10^-3^ for WT, 10^-1^ for ΔORF69) with either WT or ΔORF69. At 2 hours post-infection (hpi), the medium was replaced with 2 mL per well DMEM supplemented with 2% FCS and 0.6% methylcellulose. At 4 dpi, cells were fixed in 4% PFA for 20 min at RT and stored in PBS at 4°C until further analysis. Whole wells were imaged on a Leica DMi8 inverse widefield microscope with a THUNDER unit (Leica) and N PLAN 5x/0.12 NA objective. The setup also included 405 nm, 488 nm, 561 nm, and 640 nm laser lines and respective filter sets. The HC PL FLUOTAR L 20x/0.40 NA objective was used for higher magnification of individual plaques. For analysis, 14×14 FOV (12864 px x 12864 px) with a 10% overlap between fields of view (FOVs) were processed with a custom image processing pipeline in Fiji ImageJ (v.1.54f). A Gaussian blur (“Σ = 10”) was applied, followed by manually removing air bubbles or dust. The threshold “Otsu” (“1000-max”) was used to create a binary mask, which was then processed using “Fill Holes.” The lower size threshold for particle analysis was determined by measuring the mean size of single infected cells. Areas with a size range of >1000 µm² were considered a plaque. Results were plotted using GraphPad Prism (v.10.4.1), the two groups were tested for normality, and the Kolmogorov-Smirnov test was used to compare. For visualization of the imaging pipeline, the binary masks were color-coded using the “BAR” plugin with the “mpl-magma” LUT (range: 0-20000 µm²).

### Passaging virus and preparation of viral DNA for sequencing via Illumina Sequencing

To assess the genomic effects of MHV-68 ΔORF69 passaging on non-complementing cells, 2 × 10⁶ NIH3T3 cells were seeded in a 10 cm tissue culture dish one day before infection. Cells were infected with 1000 PFU of ΔORF69 and incubated at 37°C with 5% CO₂ until full cytopathic effect (CPE) was observed. The virus was harvested by scraping and resuspending cells in a medium, then stored at −80°C for one freeze-thaw cycle, generating passage 1 (P1). For subsequent passages, fresh NIH3T3 cells were seeded and infected with the same virus volume as the prior passage. This procedure was repeated until passage 9 (P9). For passage 10 (P10), the supernatant was collected, centrifuged at 200 rcf for 3 min to pellet debris, filtered (0.45 µm), and concentrated via centrifugation at 16,000 rcf for 2 h at 4°C. The upper supernatant was discarded, leaving 2 mL of concentrated virus. For DNA extraction, 1 mL of the virus suspension was incubated for 1 h at 50°C with 50 µg Proteinase K, 1% SDS, and 1 µL RNase A. The sample was processed using phase-lock tubes pre-spun at 20000 rcf for 2 min at 4°C. Sequential extractions with phenol-chloroform-isoamyl alcohol were performed, followed by chloroform extraction. The aqueous phase was pooled, and DNA was precipitated with sodium acetate (3 M, 1/10 volume) and isopropanol (4/5 volume), incubated at RT for 25 min, and centrifuged at 18,000 rcf for 30 min at 4°C. The DNA pellet was washed with 70% ethanol and distilled water overnight at 4°C. ΔORF69 P10 DNA was sequenced using the Illumina MiSeq platform. Reads were assembled via a reference-guided approach (reference: *in silico* MHV-68 ΔORF69 genome), and variant calling was performed.

### Sanger Sequencing

The ORFs were amplified directly from purified viral DNA (BAC, P1, P5, P10) as a template. Reactions were performed using the KOD Xtreme HotStart Polymerase (Sigma Aldrich, USA) according to the manufacturer’s instructions and primers below. PCR products were separated by 1% agarose gel electrophoresis and purified using the NucleoSpin Gel and PCR Clean-up Kit (Macherey-Nagel, Germany) according to the manufacturer’s protocol. The purified PCR fragments were sent for Sanger sequencing (ORF67, Microsynth Seqlab GmbH, Germany; ORF69 Eurofins Genomics, Germany), sequencing histograms were analyzed, and aligned to the reference sequence using SnapGene (GSL Biotech LLC, USA).

Primer Sequences:

ORF69 F: 5’-ATGCGCTCAACAGGCTCTGCT-3’

ORF69 R: 5’-TTGCTGAGAAAGACGAGATACAATGTTGAAG-3’

ORF67 F: 5’-CAGGGATACCATATTGACCCTGGTGGAC-3’

ORF67 R: 5’-GTCCAAGGCCCCCATCACCATAC-3’

ORF4 F: 5’-GTGGCCCTACCCCGAATCTC-3’

ORF4 R: 5’-GTAACCACCCACGCCGAG-3’

### Preparation of samples for Transmission Electron Microscopy

For TEM sample preparation, NIH3T3 cells were seeded in µ-Dish 35 mm, high Grid-500 (Ibidi, Germany) and infected with an MOI of 0.1 with either WT or ΔORF69. After either 1 dpi (WT) or 4 dpi ( ΔORF69), cells were fixed with 2% PFA and 2.5% GA in PBS for 20 min at RT, followed by post-fixation in 2.5% GA in PBS for 1h at 4°C. The position of infected cells was selected using a Nikon Ti2 (Nikon) spinning-disk fluorescence microscope equipped with a Yokogawa CSU-W1 SoRa spinning disc unit (Yokogawa), two Hamamatsu Orca Fusion BT sCMOS cameras, and a 20x PLAN APOλD NA=0.80 (Nikon). The setup also included 405, 488, 561, and 640 nm laser lines and respective filter sets. After the localization of cells of interest, samples were washed with PBS and incubated in 1% osmium in PBS for 20 min, then stained with uranyl acetate for 20 min in the dark. After washing, samples were dehydrated in increasing ethanol concentrations (50%, 70%, 90%, and 100%) for 10 minutes each. For embedding in epoxy resin (EPON) (89), samples were incubated in 50% EPON for 30 min, 70% EPON for 1.5h, and 100% EPON overnight. After the 100% EPON was renewed twice, samples were polymerized at 60°C overnight. Ultra-thin sections (50 nm) of multiple cells and depths were cut with a diamond knife and transferred onto copper grids. The grids were analyzed via TEM using a FEI Tecnai G20 (Thermo Fisher, USA).

### Embedding virus pellets from supernatant for TEM

For supernatant preparation for TEM, 5 × 10⁶ NIH3T3 cells were seeded on two 15 cm dishes and infected the following day with a 0.1 MOI of ΔORF69. After 1h, the medium was replaced with prewarmed DMEM containing 5% FCS. At 3 dpi, the supernatant was collected, centrifuged at 2300 rcf for 5 min, and then transferred to a new tube. The supernatant was centrifuged again, and 3 mL of 30% sucrose cushion was added to a tube, which was then centrifuged at 15,000 rcf for 1.5 h at 4°C. After removing the supernatant, the pellet was resuspended in 1 mL PBS with 1% FBS and centrifuged at 15,000 rcf for 1.5 h at 4°C. The pellet was fixed in 4% PFA in PBS and, depending on the consistency of the pellet, either directly processed for TEM or transferred into capillary tubes for support (Hollow cellulose fiber; type LD OC O2; Microdyn Wuppertal, Germany) (90). The tubes were sealed with a ring curette, processed according to the TEM preparation of cells, and analyzed by TEM using a FEI Tecnai G20 (Thermo Fisher, USA).

### Negative Stain

For negative staining of the pelleted supernatant, 10 µl PFA fixed virus suspension was applied to a piece of parafilm on ice. The glow discharge treated carbon coated grid (Science Services, Munich, Germany) was placed on a drop of virus suspension and incubated for 5min on ice. This was followed by three washing steps for 10s in double distilled water and incubation for 10s in 1% uranyl acetate in water. The grid was then dried on filter paper and analyzed by TEM using a FEI Tecnai G20 electron microscope (Thermofisher, USA).

### Quantification of capsids

Fluorescently labeled infected cells were initially identified in fluorescence microscopy images before correlating them with their corresponding regions in EM images. Individual EM images were matched to specific cells, and each image’s depth within the cell was categorized by grid number. Capsids in each image were manually scored using the “Point Tool” in Fiji/ImageJ (v.1.54f), with distinct counters assigned to different capsid types: Counter 0 = A-capsid, Counter 1 = B-capsid, Counter 2 = C-capsid (mature capsid), and Counter 3 = unclear (capsids that could not be confidently classified). The recorded counts were documented in a Microsoft Excel sheet, including metadata such as cell identifier, grid number, image number, subcellular localization (cytoplasm or nucleus), and the corresponding capsid counts for each category. This structured dataset enabled traceability of results to individual images. Total capsid numbers and ratios were calculated using Microsoft Excel, and data visualization was performed using GraphPad Prism (v.10.4.1).

### Live-cell imaging

3T3 cells were seeded in Ibidi 8-well glass bottom slides (Ibidi, Germany) at a density of 1.2 × 10⁴ cells per well. Cells were infected with an MOI of 0.1 per well with either WT or ΔORF69. At 2 hpi, the medium was replaced with 200 µL per well DMEM supplemented with 2% FCS. At 1 dpi, the nuclei were counterstained with 10µg/mL Hoechst 33342 in PBS for 15 min at 37°C. The cells were continuously imaged for 40 sec on a Nikon Ti2 Spinning disk fluorescence microscope equipped with a Yokogawa CSU-W1 SoRa spinning disc unit, two Hamamatsu Orca Fusion BT sCMOS cameras, and a 100x Apo TIRF NA=1.49 objective (Nikon). The setup also included 405 nm, 455nm, 488 nm, 561 nm, and 640 nm laser lines and respective filter sets. An environmental control system kept physiological growth conditions constant at 37°C, with 5% CO2. A representative image of the cells was color-coded based on intensity using the LUT “mpl-magma” in Fiji ImageJ (v.1.54f).

### Cytotoxicity on NIH3T3 cells

3T3 cells were seeded in 24-well plates at a density of 1.08 × 10⁵ cells per well for high concentrations (10 µM - 100µM) or 1.11 × 10⁵ cells per well for low concentrations (0.1 µM - 10µM) in 500 µL DMEM supplemented with 10% FCS and incubated overnight at 37°C, 5% CO₂. U0126 (Art.No: 662005, Sigma Aldrich, USA) and Roscovitine (Art.No: R772, Sigma Aldrich, USA) were prepared as 10 mM stock solutions in DMSO, aliquoted, and stored at -20°C. The following day, the medium was replaced with 500 µL per well of DMEM containing 2% FCS and 0.6% methylcellulose, supplemented with U0126 (10, 25, 50, or 100 µM), Roscovitine (0.1, 0.5, 1, 2.5, 5, 10, 25, 50, or 100 µM), DMSO (≤1%, vehicle control), or PBS (negative control) and incubated protected from light. The overlay was renewed daily with overlay containing the freshly diluted respective inhibitor concentrations or control solutions. At 3 dpi, cytotoxic effects were assessed by aspirating the overlay, washing cells with 500 µL PBS, and detaching them with 200 µL TrypLE Express Enzym (1x) (Gibco, ThermoFisher Scientific, USA). Trypsinization was blocked with 800 µL DMEM containing FCS, and cells were resuspended and transferred to Eppendorf tubes for counting to determine cell numbers. Values were plotted using Microsoft Excel.

### Inhibitor studies

3T3 cells were seeded in Ibidi 8-well glass bottom slides (Ibidi, Germany) at a density of 6.0 × 10⁴ cells per well for inhibitor treatment conditions and 4.5 × 10⁴ cells per well for control conditions to achieve approximately 90% and 70% confluency, respectively, the following day. Cells were infected with ∼60 PFU per well with either WT or ΔORF69. At 2 hpi, the medium was replaced with 200 µL per well DMEM supplemented with 2% FCS and 0.6% methycellulose, and either PBS (0.5%), DMSO (0.5%), Roscovitine (5 µM), or U0126 (50 µM). The overlay was renewed daily by removing as much of the existing overlay as possible and replacing it with fresh overlay containing controls or inhibitors. At 3 dpi, cells were fixed in 4% PFA for 20 min at RT and stored in PBS at 4°C until further analysis. Cells were imaged with 20×20 field of view with an ROI-size of 1024×1024 pixel with 15% overlap and stitching via blending on a Nikon Ti2 (Nikon) spinning-disk fluorescence microscope equipped with a Yokogawa CSU-W1 SoRa spinning disc unit (Yokogawa), two Hamamatsu Orca Fusion BT sCMOS cameras and a 10x PLAN APOλD NA=0.45 objective (Nikon). The setup also included 405 nm, 488 nm, 561 nm, and 640 nm laser lines and respective filter sets. FOVs were analyzed with a custom image processing pipeline in Fiji ImageJ (v.1.54f). A Gaussian blur (“Σ = 15”) was applied, followed by thresholding using “Otsu” (“110-max”) to generate a binary mask. The mask was processed with “Fill Holes” to ensure continuous regions. Plaques were identified using “Analyze Particles” with a size threshold of >1000 µm². Results were plotted using GraphPad Prism (v.10.4.1), the groups were tested for normality, and the Kruskal-Wallis test was used to compare multiple groups.

## Acknowledgments

We thank the Advanced Light and Fluorescence Microscopy (ALFM) Facility at the Centre for Structural Systems Biology (CSSB), particularly Roland Thünauer, for support with light microscopy image acquisition and analysis. We thank the Technology Platform High-Throughput Sequencing at Leibniz Institute of Virology, Hamburg, for their excellent technical support. Work in the Bosse lab was funded by the Deutsche Forschungsgemeinschaft (DFG, German Research Foundation) under Germany’s Excellence Strategy EXC 2155—project no. 390874280, the DFG-funded RTG 2771 Humans and Microbes, project no. 453548970 (Bosse), the DFG-funded RTG 2887, project number 49735088, by the Wellcome Trust through a Collaborative Award (209250/Z/17/Z) and the Leibniz ScienceCampus InterACt, funded by the BWFGB Hamburg and the Leibniz Association (W75/2022) InterACt and “Hamburg-X Infektionsforschung”. Moreover, the Bosse lab was funded through the DFG Research Unit FOR5200 DEEP-DV (443644894) project BO 4158/5-1 and the DFG CRC1648 – SFB 1648/1 2024 – 512741711”.

## Data Availability

All datasets generated and analyzed during this study are publicly available via the Zenodo repository. The dataset includes microscopy data, comprising files from electron microscopy, live-cell imaging, and conventional light microscopy, as well as the gel image of the BAC restriction digest. Sequencing data, including next-generation sequencing (NGS) and Sanger sequencing experiments. Protein structure predictions were generated using AlphaFold.

The complete dataset can be accessed via Zenodo using the following DOI: 10.5281/zenodo.15147695.

## References

1. Knipe DM, Howley PM. Fields’ Virology. Lippincott Williams & Wilkins; 2007. 3116 p.

2. Rajčáni J, Kúdelová M. Murine Herpesvirus Pathogenesis: A Model for the Analysis of Molecular Mechanisms of Human Gamma Herpesvirus Infections. 2005 Jul 22 [cited 2025 Mar 3]; Available from: https://akjournals.com/view/journals/030/52/1/article-p41.xml

3. Heming JD, Conway JF, Homa FL. Herpesvirus Capsid Assembly and DNA Packaging. Adv Anat Embryol Cell Biol. 2017;223:119–42.

4. Sherman G, Bachenheimer SL. Characterization of intranuclear capsids made by ts morphogenic mutants of HSV-1. Virology. 1988 Apr 1;163(2):471–80.

5. Lösing J, Häge S, Schütz M, Wagner S, Wardin J, Sticht H, et al. “Shared-Hook” and “Changed-Hook” Binding Activities of Herpesviral Core Nuclear Egress Complexes Identified by Random Mutagenesis. Cells. 2022 Dec 13;11(24):4030.

6. Klupp BG, Mettenleiter TC. The Knowns and Unknowns of Herpesvirus Nuclear Egress. Annu Rev Virol. 2023 Sep 29;10(1):305–23.

7. Lötzerich M, Ruzsics Z, Koszinowski UH. Functional Domains of Murine Cytomegalovirus Nuclear Egress Protein M53/p38. Journal of Virology. 2006 Jan;80(1):73–84.

8. Bubeck A, Wagner M, Ruzsics Z, Lötzerich M, Iglesias M, Singh IR, et al. Comprehensive Mutational Analysis of a Herpesvirus Gene in the Viral Genome Context Reveals a Region Essential for Virus Replication. Journal of Virology. 2004 Aug;78(15):8026–35.

9. Marschall M, Häge S, Conrad M, Alkhashrom S, Kicuntod J, Schweininger J, et al. Nuclear Egress Complexes of HCMV and Other Herpesviruses: Solving the Puzzle of Sequence Coevolution, Conserved Structures and Subfamily-Spanning Binding Properties. Viruses. 2020 Jun 24;12(6):683.

10. Schweininger J, Kriegel M, Häge S, Conrad M, Alkhashrom S, Lösing J, et al. The crystal structure of the varicella-zoster Orf24-Orf27 nuclear egress complex spotlights multiple determinants of herpesvirus subfamily specificity. Journal of Biological Chemistry [Internet]. 2022 Mar 1 [cited 2025 Mar 3];298(3). Available from: https://www.jbc.org/article/S0021-9258(22)00065-5/abstract

11. Thorsen MK, Draganova EB, Heldwein EE. The nuclear egress complex of Epstein-Barr virus buds membranes through an oligomerization-driven mechanism. PLoS Pathog. 2022 Jul;18(7):e1010623.

12. Bigalke JM, Heldwein EE. Structural basis of membrane budding by the nuclear egress complex of herpesviruses. EMBO J. 2015 Dec 2;34(23):2921–36.

13. Lye MF, Sharma M, El Omari K, Filman DJ, Schuermann JP, Hogle JM, et al. Unexpected features and mechanism of heterodimer formation of a herpesvirus nuclear egress complex. EMBO J. 2015 Dec 2;34(23):2937–52.

14. Muller YA, Häge S, Alkhashrom S, Höllriegl T, Weigert S, Dolles S, et al. High-resolution crystal structures of two prototypical β- and γ-herpesviral nuclear egress complexes unravel the determinants of subfamily specificity. J Biol Chem. 2020 Mar 6;295(10):3189–201.

15. Walzer SA, Egerer-Sieber C, Sticht H, Sevvana M, Hohl K, Milbradt J, et al. Crystal Structure of the Human Cytomegalovirus pUL50-pUL53 Core Nuclear Egress Complex Provides Insight into a Unique Assembly Scaffold for Virus-Host Protein Interactions. J Biol Chem. 2015 Nov 13;290(46):27452–8.

16. Zeev-Ben-Mordehai T, Weberruß M, Lorenz M, Cheleski J, Hellberg T, Whittle C, et al. Crystal Structure of the Herpesvirus Nuclear Egress Complex Provides Insights into Inner Nuclear Membrane Remodeling. Cell Rep. 2015 Dec 29;13(12):2645–52.

17. Hagen C, Dent KC, Zeev-Ben-Mordehai T, Grange M, Bosse JB, Whittle C, et al. Structural Basis of Vesicle Formation at the Inner Nuclear Membrane. Cell. 2015 Dec 17;163(7):1692–701.

18. Pražák V, Mironova Y, Vasishtan D, Hagen C, Laugks U, Jensen Y, et al. Molecular plasticity of herpesvirus nuclear egress analysed in situ. Nat Microbiol. 2024 Jun 25;9(7):1842–55.

19. Bigalke JM, Heuser T, Nicastro D, Heldwein EE. Membrane deformation and scission by the HSV-1 nuclear egress complex. Nat Commun. 2014 Jun 11;5(1):4131.

20. Jumper J, Evans R, Pritzel A, Green T, Figurnov M, Ronneberger O, et al. Highly accurate protein structure prediction with AlphaFold. Nature. 2021;596(7873):583–9.

21. Akdel M, Pires DEV, Pardo EP, Jänes J, Zalevsky AO, Mészáros B, et al. A structural biology community assessment of AlphaFold2 applications. Nat Struct Mol Biol. 2022;29(11):1056–67.

22. Baek M, DiMaio F, Anishchenko I, Dauparas J, Ovchinnikov S, Lee GR, et al. Accurate prediction of protein structures and interactions using a 3-track neural network. Science. 2021 Aug 20;373(6557):871–6.

23. Soh TK, Ognibene S, Sanders S, Schäper R, Kaufer BB, Bosse JB. A proteome-wide structural systems approach reveals insights into protein families of all human herpesviruses. Nat Commun. 2024 Nov 26;15(1):10230.

24. Nomburg J, Doherty EE, Price N, Bellieny-Rabelo D, Zhu YK, Doudna JA. Birth of protein folds and functions in the virome. Nature. 2024 Sep;633(8030):710–7.

25. Mifsud JCO, Lytras S, Oliver MR, Toon K, Costa VA, Holmes EC, et al. Mapping glycoprotein structure reveals Flaviviridae evolutionary history. Nature. 2024 Sep;633(8030):695–703.

26. Wild P, Senn C, Manera CL, Sutter E, Schraner EM, Tobler K, et al. Exploring the Nuclear Envelope of Herpes Simplex Virus 1-Infected Cells by High-Resolution Microscopy. Journal of Virology. 2009 Jan;83(1):408–19.

27. Leuzinger H, Ziegler U, Schraner EM, Fraefel C, Glauser DL, Heid I, et al. Herpes simplex virus 1 envelopment follows two diverse pathways. J Virol. 2005 Oct;79(20):13047–59.

28. Farina A, Feederle R, Raffa S, Gonnella R, Santarelli R, Frati L, et al. BFRF1 of Epstein-Barr virus is essential for efficient primary viral envelopment and egress. J Virol. 2005 Mar;79(6):3703–12.

29. Gonnella R, Farina A, Santarelli R, Raffa S, Feederle R, Bei R, et al. Characterization and Intracellular Localization of the Epstein-Barr Virus Protein BFLF2: Interactions with BFRF1 and with the Nuclear Lamina. Journal of Virology. 2005 Mar 15;79(6):3713–27.

30. Santarelli R, Farina A, Granato M, Gonnella R, Raffa S, Leone L, et al. Identification and characterization of the product encoded by ORF69 of Kaposi’s sarcoma-associated herpesvirus. J Virol. 2008 May;82(9):4562–72.

31. Lv Y, Shen S, Xiang L, Jia X, Hou Y, Wang D, et al. Functional Identification and Characterization of the Nuclear Egress Complex of a Gammaherpesvirus. J Virol. 2019 Dec 15;93(24):e01422–19.

32. Virgin HW, Latreille P, Wamsley P, Hallsworth K, Weck KE, Dal Canto AJ, et al. Complete sequence and genomic analysis of murine gammaherpesvirus 68. J Virol. 1997 Aug;71(8):5894– 904.

33. Desai PJ, Pryce EN, Henson BW, Luitweiler EM, Cothran J. Reconstitution of the Kaposi’s sarcoma-associated herpesvirus nuclear egress complex and formation of nuclear membrane vesicles by coexpression of ORF67 and ORF69 gene products. Journal of virology. 2012;86(1):594–8.

34. Lv Y, Shen S, Xiang L, Jia X, Hou Y, Wang D, et al. Functional Identification and Characterization of the Nuclear Egress Complex of a Gammaherpesvirus. Journal Of Virology. 2019;93(24).

35. Bosse JB, Virding S, Thiberge SY, Scherer J, Wodrich H, Ruzsics Z, et al. Nuclear herpesvirus capsid motility is not dependent on F-actin. MBio. 2014;5(5).

36. Sherry MR, Hay TJM, Gulak MA, Nassiri A, Finnen RL, Banfield BW. The Herpesvirus Nuclear Egress Complex Component, UL31, Can Be Recruited to Sites of DNA Damage Through Poly-ADP Ribose Binding. Sci Rep. 2017 May 15;7(1):1882.

37. Thorsen MK, Lai A, Lee MW, Hoogerheide DP, Wong GCL, Freed JH, et al. Highly Basic Clusters in the Herpes Simplex Virus 1 Nuclear Egress Complex Drive Membrane Budding by Inducing Lipid Ordering. mBio. 2021 Aug 24;12(4):10.1128/mbio.01548-21.

38. Abramson J, Adler J, Dunger J, Evans R, Green T, Pritzel A, et al. Accurate structure prediction of biomolecular interactions with AlphaFold 3. Nature. 2024 Jun;630(8016):493–500.

39. Häge S, Horsch D, Stilp AC, Kicuntod J, Müller R, Hamilton ST, et al. A quantitative nuclear egress assay to investigate the nucleocytoplasmic capsid release of human cytomegalovirus. Journal of Virological Methods. 2020 Sep 1;283:113909.

40. Dai YC, Liao YT, Juan YT, Cheng YY, Su MT, Su YZ, et al. The Novel Nuclear Targeting and BFRF1-Interacting Domains of BFLF2 Are Essential for Efficient Epstein-Barr Virus Virion Release. J Virol. 2020 Jan 17;94(3):e01498–19.

41. Klupp B, Granzow H, Mettenleiter T. Nuclear Envelope Breakdown Can Substitute for Primary Envelopment-Mediated Nuclear Egress of Herpesviruses. Journal Of Virology. 2011;85(16):8285– 92.

42. Grimm K, Klupp B, Granzow H, Muller F, Fuchs W, Mettenleiter T. Analysis of Viral and Cellular Factors Influencing Herpesvirus-Induced Nuclear Envelope Breakdown. Journal Of Virology. 2012;86(12):6512–21.

43. Clambey ET, Virgin HW, Speck SH. Characterization of a spontaneous 9.5-kilobase-deletion mutant of murine gammaherpesvirus 68 reveals tissue-specific genetic requirements for latency. Journal of virology. 2002;76(13):6532–44.

44. Oda W, Mistrikova J, Stancekova M, Dutia BM, Nash AA, Takahata H, et al. Analysis of genomic homology of murine gammaherpesvirus (MHV)-72 to MHV-68 and impact of MHV-72 on the survival and tumorigenesis in the MHV-72-infected CB17 scid/scid and CB17+/+ mice. Pathology International. 2005;55(9):558–68.

45. Macrae AI, Dutia BM, Milligan S, Brownstein DG, Allen DJ, Mistrikova J, et al. Analysis of a novel strain of murine gammaherpesvirus reveals a genomic locus important for acute pathogenesis. J Virol. 2001 Jun;75(11):5315–27.

46. Chalupkova A, Hricová M, Hrabovska Z, Mistríková J. Pathogenetical Characterization of MHV76: a Spontaneous 9.5-Kilobase-Deletion Mutant of Murine Lymphotropic Gammaherpesvirus 68. Acta Veterinaria Brno - ACTA VET BRNO. 2008 Jun 1;77:231–7.

47. Grimm KS, Klupp BG, Granzow H, Müller FM, Fuchs W, Mettenleiter TC. Analysis of viral and cellular factors influencing herpesvirus-induced nuclear envelope breakdown. J Virol. 2012 Jun;86(12):6512–21.

48. Chang YE, Van Sant C, Krug PW, Sears AE, Roizman B. The null mutant of the U(L)31 gene of herpes simplex virus 1: construction and phenotype in infected cells. J Virol. 1997 Nov;71(11):8307–15.

49. Popa M, Ruzsics Z, Lötzerich M, Dölken L, Buser C, Walther P, et al. Dominant Negative Mutants of the Murine Cytomegalovirus M53 Gene Block Nuclear Egress and Inhibit Capsid Maturation. J Virol. 2010 Sep;84(18):9035–46.

50. Granato M, Feederle R, Farina A, Gonnella R, Santarelli R, Hub B, et al. Deletion of Epstein-Barr Virus BFLF2 Leads to Impaired Viral DNA Packaging and Primary Egress as Well as to the Production of Defective Viral Particles. Journal of Virology. 2008 Apr 15;82(8):4042–51.

51. Schulz KS, Klupp BG, Granzow H, Mettenleiter TC. Glycoproteins gB and gH Are Required for Syncytium Formation but Not for Herpesvirus-Induced Nuclear Envelope Breakdown. Journal of Virology. 2013 Sep;87(17):9733–41.

52. Maric M, Haugo AC, Dauer W, Johnson DC, Roller R. Nuclear envelope breakdown induced by herpes simplex virus type 1 involves the activity of viral fusion proteins. Virology. 2014;460– 461:128–37.

53. Desai PJ, Pryce EN, Henson BW, Luitweiler EM, Cothran J. Reconstitution of the Kaposi’s Sarcoma-Associated Herpesvirus Nuclear Egress Complex and Formation of Nuclear Membrane Vesicles by Coexpression of ORF67 and ORF69 Gene Products. Journal of Virology. 2012 Jan;86(1):594–8.

54. Klupp B, Granzow H, Fuchs W, Keil G, Finke S, Mettenleiter T. Vesicle formation from the nuclear membrane is induced by coexpression of two conserved herpesvirus proteins. Proceedings Of The National Academy Of Sciences. 2007;104(17):7241–6.

55. Tandon R, Mocarski ES. Viral and host control of cytomegalovirus maturation. Trends in Microbiology. 2012 Aug 1;20(8):392–401.

56. Hogue IB. Tegument Assembly, Secondary Envelopment and Exocytosis. Current Issues in Molecular Biology. 2021 Mar;42(1):551–604.

57. Meijer L, Borgne A, Mulner O, Chong JP, Blow JJ, Inagaki N, et al. Biochemical and cellular effects of roscovitine, a potent and selective inhibitor of the cyclin-dependent kinases cdc2, cdk2 and cdk5. Eur J Biochem. 1997 Jan 15;243(1–2):527–36.

58. Stahl JA, Chavan SS, Sifford JM, MacLeod V, Voth DE, Edmondson RD, et al. Phosphoproteomic analyses reveal signaling pathways that facilitate lytic gammaherpesvirus replication. PLoS Pathog. 2013 Sep;9(9):e1003583.

59. Duncia JV, Santella JB, Higley CA, Pitts WJ, Wityak J, Frietze WE, et al. MEK inhibitors: The chemistry and biological activity of U0126, its analogs, and cyclization products. Bioorganic & Medicinal Chemistry Letters. 1998 Oct 20;8(20):2839–44.

60. Luitweiler E, Henson B, Pryce E, Patel V, Coombs G, McCaffery J, et al. Interactions of the Kaposi’s Sarcoma-Associated Herpesvirus Nuclear Egress Complex: ORF69 Is a Potent Factor for Remodeling Cellular Membranes. Journal Of Virology. 2013;87(7):3915–29.

61. Hellberg T, Paßvogel L, Schulz KS, Klupp BG, Mettenleiter TC. Chapter Three - Nuclear Egress of Herpesviruses: The Prototypic Vesicular Nucleocytoplasmic Transport. In: Kielian M, Maramorosch K, Mettenleiter TC, editors. Advances in Virus Research [Internet]. Academic Press; 2016 [cited 2025 Mar 13]. p. 81–140. Available from: https://www.sciencedirect.com/science/article/pii/S0065352715000913

62. Fuchs W, Klupp BG, Granzow H, Osterrieder N, Mettenleiter TC. The Interacting UL31 and UL34 Gene Products of Pseudorabies Virus Are Involved in Egress from the Host-Cell Nucleus and Represent Components of Primary Enveloped but Not Mature Virions. Journal of Virology. 2002 Jan;76(1):364–78.

63. Reynolds AE, Ryckman BJ, Baines JD, Zhou Y, Liang L, Roller RJ. UL31 and UL34 Proteins of Herpes Simplex Virus Type 1 Form a Complex That Accumulates at the Nuclear Rim and Is Required for Envelopment of Nucleocapsids. Journal of Virology. 2001 Sep 15;75(18):8803–17.

64. Sharma M, Kamil JP, Coughlin M, Reim NI, Coen DM. Human Cytomegalovirus UL50 and UL53 Recruit Viral Protein Kinase UL97, Not Protein Kinase C, for Disruption of Nuclear Lamina and Nuclear Egress in Infected Cells. J Virol. 2014 Jan;88(1):249–62.

65. Klupp BG, Granzow H, Mettenleiter TC. Primary Envelopment of Pseudorabies Virus at the Nuclear Membrane Requires the UL34 Gene Product. Journal of Virology. 2000 Nov;74(21):10063–73.

66. Roller RJ, Zhou Y, Schnetzer R, Ferguson J, DeSalvo D. Herpes Simplex Virus Type 1 UL34 Gene Product Is Required for Viral Envelopment. Journal of Virology. 2000 Jan;74(1):117–29.

67. Häge S, Sonntag E, Svrlanska A, Borst E, Stilp AC, Horsch D, et al. Phenotypical Characterization of the Nuclear Egress of Recombinant Cytomegaloviruses Reveals Defective Replication upon ORF-UL50 Deletion but Not pUL50 Phosphosite Mutation. Viruses. 2021 Jan 22;13:165.

68. Ye GJ, Roizman B. The essential protein encoded by the UL31 gene of herpes simplex virus 1 depends for its stability on the presence of UL34 protein. Proceedings of the National Academy of Sciences. 2000 Sep 26;97(20):11002–7.

69. Newcomb WW, Homa FL, Brown JC. Herpes Simplex Virus Capsid Structure: DNA Packaging Protein UL25 Is Located on the External Surface of the Capsid near the Vertices. Journal of Virology. 2006 Jul;80(13):6286–94.

70. Tandon R, Mocarski ES, Conway JF. The A, B, Cs of Herpesvirus Capsids. Viruses. 2015 Mar;7(3):899–914.

71. Klupp BG, Granzow H, Keil GM, Mettenleiter TC. The capsid-associated UL25 protein of the alphaherpesvirus pseudorabies virus is nonessential for cleavage and encapsidation of genomic DNA but is required for nuclear egress of capsids. J Virol. 2006 Jul;80(13):6235–46.

72. Yang K, Wills E, Lim HY, Zhou ZH, Baines JD. Association of herpes simplex virus pUL31 with capsid vertices and components of the capsid vertex-specific complex. J Virol. 2014 Apr;88(7):3815–25.

73. Takeshima K, Arii J, Maruzuru Y, Koyanagi N, Kato A, Kawaguchi Y. Identification of the Capsid Binding Site in the Herpes Simplex Virus 1 Nuclear Egress Complex and Its Role in Viral Primary Envelopment and Replication. J Virol. 2019 Oct 15;93(21):e01290–19.

74. Yang K, Baines JD. Selection of HSV capsids for envelopment involves interaction between capsid surface components pUL31, pUL17, and pUL25. Proc Natl Acad Sci U S A. 2011 Aug 23;108(34):14276–81.

75. Bailer S. Venture from the Interior—Herpesvirus pUL31 Escorts Capsids from Nucleoplasmic Replication Compartments to Sites of Primary Envelopment at the Inner Nuclear Membrane. Cells. 2017;6:null.

76. Leelawong M, Guo D, Smith GA. A physical link between the pseudorabies virus capsid and the nuclear egress complex. Journal of virology. 2011;85(22):11675–84.

77. Schnee M, Ruzsics Z, Bubeck A, Koszinowski UH. Common and Specific Properties of Herpesvirus UL34/UL31 Protein Family Members Revealed by Protein ComplementationAssay. Journal of Virology. 2006 Dec;80(23):11658–66.

78. Speese SD, Ashley J, Jokhi V, Nunnari J, Barria R, Li Y, et al. Nuclear envelope budding enables large ribonucleoprotein particle export during synaptic Wnt signaling. Cell. 2012 May 11;149(4):832–46.

79. Zacny VL, Wilson J, Pagano JS. The Epstein-Barr virus immediate-early gene product, BRLF1, interacts with the retinoblastoma protein during the viral lytic cycle. J Virol. 1998 Oct;72(10):8043–51.

80. Izumiya Y, Lin SF, Ellison TJ, Levy AM, Mayeur GL, Izumiya C, et al. Cell cycle regulation by Kaposi’s sarcoma-associated herpesvirus K-bZIP: direct interaction with cyclin-CDK2 and induction of G1 growth arrest. Journal of virology. 2003;77(17):9652–61.

81. Upton JW, van Dyk LF, Speck SH. Characterization of murine gammaherpesvirus 68 v-cyclin interactions with cellular cdks. Virology. 2005 Oct 25;341(2):271–83.

82. Tishchenko A, Romero N, Waesberghe CV, Delva JL, Vickman O, Smith GA, et al. Pseudorabies virus infection triggers pUL46-mediated phosphorylation of connexin-43 and closure of gap junctions to promote intercellular virus spread. PLOS Pathogens. 2025 Jan 21;21(1):e1012895.

83. Adler H, Messerle M, Wagner M, Koszinowski UH. Cloning and Mutagenesis of the Murine Gammaherpesvirus 68 Genome as an Infectious Bacterial Artificial Chromosome. Journal of Virology. 2000 Aug;74(15):6964–74.

84. Vidy A, Sacher T, Adler H, Jordan S, Koszinowski UH, Ruzsics Z. Systemic and Local Infection Routes Govern Different Cellular Dissemination Pathways during Gammaherpesvirus Infection In Vivo. Journal of Virology. 2013 Apr 15;87(8):4596–608.

85. Bosse JB. Herpesvirus capsid dynamics in living cells [Internet] [Text.PhDThesis]. Ludwig-Maximilians-Universität München; 2011 [cited 2025 Mar 18]. Available from: https://edoc.ub.uni-muenchen.de/14219/

86. Cherepanov PP, Wackernagel W. Gene disruption in *Escherichia coli*: TcR and KmR cassettes with the option of Flp-catalyzed excision of the antibiotic-resistance determinant. Gene. 1995 Jan 1;158(1):9–14.

87. Bosse JB, Bauerfeind R, Popilka L, Marcinowski L, Taeglich M, Jung C, et al. A beta-herpesvirus with fluorescent capsids to study transport in living cells. PLoS One. 2012;7(7):e40585.

88. Tischer BK, Smith GA, Osterrieder N. En passant mutagenesis: a two step markerless red recombination system. Methods Mol Biol. 2010;634:421–30.

89. Luft JH. Improvements in epoxy resin embedding methods. The Journal of biophysical and biochemical cytology. 1961;9(2):409.

90. Hohenberg H, Mannweiler K, Müller M. High-pressure freezing of cell suspensions in cellulose capillary tubes. Journal of microscopy. 1994;175(1):34–43.

